# FastGA: Fast Genome Alignment

**DOI:** 10.1101/2025.06.15.659750

**Authors:** Gene Myers, Richard Durbin, Chenxi Zhou

## Abstract

FastGA finds alignments between two genome sequences more than an order of magnitude faster than previous methods that have comparable sensitivity. Its speed is due to (a) a carefully engineered architecture involving only *cache-coherent MSD radix sorts* and merges, (b) a novel algorithm for finding *adaptive seed* hits in a linear merge of sorted k-mer tables, and (c) a variant of the Myers *adaptive wave* algorithm [1] to find alignments around a chain of seed hits that detects alignments with up to 25-30% variation. It further does not require pre-masking of repetitive sequence, and stores millions of alignments in a fraction of the space of a conventional CIGAR-string [2] using a *trace-point* encoding that is further compressed by the ONEcode data system [3] introduced here.

As an example, two bat genomes of size 2.2Gbp and 2.5Gbp can be compared in a little over 2 minutes using 8 threads on an Apple M4 Max laptop using 5.7GB of memory and producing 1.05 million alignments totaling 1.63Gbp of aligned sequence that cover about 60% of each genome. The output “ALN”-formatted file occupies 66MB. This file can be converted to a PAF file with CIGAR strings in 6 seconds, where the PAF representation is a significantly larger 1.03GB file.

FastGA is freely available at github: http://www.github.com/thegenemyers/FASTGA along with utilities for viewing inputs, intermediate files, and outputs and transforming outputs into other common formats. Specifically, FastGA can, in addition to its highly efficient ONEcode representation, output PSL-formatted alignments, or PAF-formatted alignments with or without CIGAR strings explicitly encoding the alignments. There is also a utility to chain FastGA’s alignments and display them in a dot-plot like view in Postscript files, and an interactive viewer is in development.

## 1 Introduction & Summary

Initiatives such as the Vertebrate Genomes Project (VGP), the Darwin Tree of Life project (DToL), and the Earth Biogenome Project (EBP) portend the *de novo* sequencing of tens of thousands of species to a near perfect standard [4, 5, 6] using single-molecule long-read sequencing technologies[7]. The reconstructions typically have chromosomal scaffolds with tens to a few hundred gaps between contigs and a consensus error rate of better than 10^−5^ [8]. The number of genomes being so sequenced is increasing geometrically, and includes species with large genomes, 10Gbp or greater. Already as of this writing, several thousand such genomes have been produced, some concentrated on particular clades, for example, over 100 of the 1400 extant bat species have been sequenced [9]. We therefore anticipate that comparing such clade-based genome collections to understand evolution at the genetic and molecular level will be a major research thrust going forward. A central computational task for this endeavor is finding all statistically significant local alignments between two complete genome sequences. Achieving this computation efficiently and accurately is thus of importance. This paper presents a suite of engineering concepts and algorithms resulting in a software tool, FastGA, that is typically an order of magnitude faster than the current fastest tools, and almost two orders of magnitude faster than the most accurate of the current tools while delivering the same sensitivity.

Almost every efficient biosequence similarity search method involves a filter that limits the search to smaller pairs of regions to be aligned, followed by a deterministic alignment algorithm that verifies whether or not a good alignment is present. For example, Blast’s seed-and-extend strategy was an early example of this [10] and the five precursors to FastGA that we discuss here follow the same overarching design with the differences being in the methods for filtration and verification.

We carefully separate the problems of genome alignment and genome homology. The former is the problem of finding all regions that have a statistically significant alignment between them, and the latter is the problem of finding regions that evolved from the same region of an ancestral genome. Not all homologous sequences will have a statistically significant alignment, because there may have been too much mutation since the last common ancestor. On the other hand, repetitive sequences, while they may share a common ancestor in themselves due to deriving from the same mobile element, can be embedded in unrelated regions of a genome, making them misleading indicators of regional genome homology. In this paper we focus on the first problem of finding statistically significant alignment. FastGA finds pairs of segments that significantly align essentially across their whole sequences, with maximal gap sizes of approximately 40bp. It does not itself chain these alignments together across larger gaps, which would suggest homology across the intervening sequence, but rather leaves this to a second step. We provide a simple chaining tool as an initial step towards the second step, but also refer users interested in larger scale homology and genome evolution to existing tools that can take alignments as input, such as the multiple genome aligner Cactus [11, 12]. We note that by not conflating these two steps, FastGA can be used for other downstream tasks such as finding recurrent insertions due to transposable elements[13].

One of the earlier genome alignment tools, and as our empirical results show still the most sensitive, is LastZ ([14, 15, 16]) developed by Webb Miller and his Ph.D. Student Robert Harris in the mid-2000’s when the only large genomes of a decent quality to compare were human and mouse. It compares every pair of assembly contigs using a Blast-like seed and extend strategy entailing zone-based dynamic programming to find alignments with a scoring function that favors transitions over transversions. It uses spaced seeds [17], where the seed pattern is user-specifiable, and provides a method for inferring an “optimal” alignment scoring function based on the overall similarity of the input genomes.

NUCmer ([18]) is another early but still frequently used aligner, whose underlying machinery is based on suffix arrays and the idea of maximal unique matches or MUMs as the seed hits. With suffix arrays, MUMs can be found in linear time and seed uniquely corresponding regions between genomes. NUCmer is part of an overall package MUMmer that predates LastZ and was developed primarily for aligning the bacterial genomes that were available at the time, but it is nonetheless general purpose. The suffix-array based seeding strategy is relatively more efficient than LastZ and simpler post-chaining alignment verification give it better time performance albeit it is less sensitive.

Minimap2 ([19, 20]) is a more recent general-purpose DNA alignment approach that was originally developed with the goal of mapping reads to genomes. The key idea is to reduce the number of k-mers inspected for seed matches by using only those that are minimizers in a window of some small size [21]. Another notable feature is the use of affine gaps often of a very large size. Using such a gap function, a collinear run of alignments that FastGA would report individually are bundled into a single macro-alignment. This makes the results look less fragmented, but from our perspective conflates the detection of significant alignments with inference from them about regional homology, and once bundled it is cumbersome to have to consider the possibility that a reported synteny block is not homologous.

Wfmash ([22])is a more recent arrival, having been designed in the context of producing a pangenome graph of the human genome. Unlike other aligners, it uses a Min-hash estimate of the Jaccard similarity between two regions [23] as the filter for finding potentially similar regions and then a recent affine gap wave-alignment algorithm ([24]) for verification. The mash-map produced by the filter requires larger regions of similarity and higher levels of identity than k-mer seed approaches as reflected in the results of the empirical experiments in this paper. But it is faster to compute and sufficient when species are proximal, such as for example a human to human or human to gorilla comparison. The use of an affine wave as opposed to a unit cost wave does not increase sensitivity but does produce nicer looking alignments as we will show in Subsection 3.2.2. It does however increase time several fold as it must compute a number of waves equal to the cost of the first gap. We advocate in this work that one should employ an affine gap penalty only as a means of refining found alignments for presentation to users.

FastGA attains its performance gains over these previous methods through both engineering and algorithmic innovations. The paper first covers the engineering ideas, specifically the importance of cache coherence and the repeated use of most-significant-digit (MSD) radix sorting to help achieve this. We use these to (1) build our version of a genome index, (2) find seed hits, and (3) find chains of seeds. The next section then treats the several algorithmic innovations in FastGA: (1) a linear index merge to find all adaptive seeds, (2) the idea of tracepoints for the space efficient representation of alignments, and (3) alignment refinement under an affine gap model. A brief section then presents the software architecture of FastGA and its underpinning in ONEcode, a general data handling framework. We conclude with an empirical comparison of the performance of FastGA versus LastZ, NUCmer, Minimap2, and wfmash under a number of scenarios. This last section importantly supports our claims of speed and sensitivity.

### 1.1 String Formalisms: Syncmers and Adaptamers

All strings *A* = *a*_1_*a*_2_ … *a*_*n*_ are assumed to be over the DNA alphabet Σ = {*a, c, g, t*}. Position *i* in *A* is the location *between* characters *a*_*i*_ and *a*_*i*+1_, so that the substring *A*[*i, j*] = *a*_*i*+1_*a*_*i*+2_ … *a*_*j*_ and is of length *j* − *i*. A *k*-mer is any substring of length *k*. The *complement* of a DNA string is *A*^*c*^ = *c*(*a*_*n*_)*c*(*a*_*n*−1_) … *c*(*a*_1_) where *c* : [*acgt*] → [*tgca*]. We will assume throughout that there is a function *φ* that assigns an integer value to any string and hence orders them. The *canonical value* Φ(*α*) of a string *α* is the smaller of the value of *α* or its complement, i.e. Φ(*α*) = *min*(*φ*(*α*), *φ*(*α*^*c*^)).

Much work has focused on *minimizers* as a representative subset of the *k*-mers of a string [21]. In this work we will use the *syncmer* concept of Bob Edgar [25]. Given *k*-mer *α* and *m < k, α* is a *closed* (*k, m*) *syncmer* if and only if the canonical value of the *m*-mer at the start or end of *α* is the smallest of all *m*-mers of *α*, i.e. *min*(Φ(*α*[0, *m*]), Φ(*α*[*k* −*m, k*])) = *min*_*i*∈[0,*k*−*m*]_ Φ(*α*[*i, i* + *m*]). Edgar showed that the subsequence of *k*-mers of a long string *A* that are closed (*k, m*) syncmers are at most *k*−*m* positions apart and on average are slightly more than (*k* − *m*)*/*2 from each other. Thus a series of closed syncmers represent a sub-sampling of the *k*-mers of a long sequence with an average density of approximately 2*/*(*k* − *m*) that are further guaranteed to cover the sequence.

Most algorithms that search for approximate string matches use the concept of *seed hits* to filter the space of possible corresponding intervals between the two strings. The most basic of these is, for some *k*, to find pairs of positions for which the *k*-mers beginning at them are the same. Immediately there is the question of how to choose *k*: too small and there will be an overwhelming, quadratically growing number of seed pairs, too large and approximate matches will be missed. That is, there is a sensitivity:specificity trade-off that depends critically on the degree of similarity sought in a match and the size and repetitiveness of the underlying sequences. One interesting solution is to seek only the *maximally unique matches*, or *MUMs* [18], that is, a sequence that begins at a pair of positions in the two sequences that is of maximal length over all such sequences. While generally selective of matching regions it may not identify multiple similar regions. A related idea that is less stringent is that of *adaptive seeds* [26] that are locally maximal as opposed to globally maximal. We call such seeds *adaptamers* throughout this paper. At a given position *i* in a sequence, the adaptamer at position *i* is the longest string beginning there, that can also be found in the other sequence or its complement. Unlike MUMs, the concept is not symmetric, i.e., if *A*[*i, i* + *p*] is an adaptamer at *i* and it matches say*B*[*j, j* + *p*], it is not necessarily the case that *B*[*j, j* + *p*] is the adaptamer at *j* but a prefix of said. Given a threshold *τ* > 0, we will deem an adaptamer *repetitive* if it or its complement occurs more than *τ* times in the other sequence.

## 2 Engineering

### 2.1 Cache Coherent Sorting for Speed

CPUs are designed to operate at the fastest clock rate possible, and memories are designed to provide the maximum density/capacity possible, at lower random access rates. So as computer hardware has advanced over the last several decades, the mismatch between the speed of the CPU and primary memory has increased, to the point where today memory latency is typically about 100 times the clock rate of the CPU. To alleviate this mismatch, multiple layers of cache, becoming increasing smaller and faster closer to the CPU, have been introduced. But it is still the case that a program that makes random accesses to a large data structure, e.g. a hash table of all 40-mers of a genome, suffers a cache miss at every layer almost every time a hash add or lookup is performed. On the other hand, *cache-coherent* programs that make a limited number of linear sweeps through memory at the same time, either increasing or decreasing, tend to suffer almost no memory latency as the cache hardware operates in multi-word blocks and can anticipate and preload blocks for such simple access patterns.

Let the notation *κ*_*i*_ refer to the *k*-mer that begins at position *i* in a sequence. A typical comparison between two genomes, *G* and *H*, would have the outline:

> Create a *k*-mer index ℋ of *H* (e.g. suffix tree, hash table, FM-index)
>
> **For** each position *p* in *G* in order **do**
>
> Look up *κ*_*p*_ in ℋ and for all positions *q* found, report (*p, q*)

There is almost no coherence between where the data for *κ*_*i*_ and *κ*_*i*+1_ are located in the memory of ℋ (at least for all the data structures listed above) so the lookup will almost always entail a cache miss through all layers of cache if the index is large enough to index a gigabase-scale genome. Instead consider using a highly optimized cache coherent sort such as a Most-Significant Digit (MSD) radix sort[27] in the following outline:

> Build a list 𝒜 = {(*p, κ*_*p*_)} of each position and its k-mer in *G* and MSD sort on the *k*-mer
>
> Build a list ℬ = {(*q, κ*_*q*_)} of each position and its k-mer in *H* and MSD sort on the *k*-mer
>
> Merge 𝒜 and ℬ in order of *k*-mer and make a list *C* of pairs (*p, q*) with the same *k*-mer
>
> Sort 𝒞 in lexicographical order of *p* and then *q* with an MSD sort

First observe that the entire computation is cache coherent. Second, the final sorted list 𝒞 is exactly the sequence of pairs output by the first approach. So in essence both methods produce the set of all seed hits, that is, pairs of locations in the underlying dynamic programming matrix that share the same *k*-mer. Naively, the first method might seem preferable as it involves fewer operations (*O*(*N*) not *O*(*N* log *N*)) and potentially less space, as the hits can be clustered and processed as they are discovered. But while the second approach excutes more operations and utilizes more space, there are no cache misses, so the average execution time of each instruction is much closer to that of the CPU clock. Moreover the space occupied by 𝒞 is of the same order as the index ℋ of the first approach if one takes some care to avoid highly repetitive *k*-mers. As the empirical results of this paper will show, FastGA, which uses the sorting approach is typically an order of magnitude or more faster than other contemporary tools all of which use a form of the first, cache-incoherent approach. The speed of FastGA makes the point that algorithms/software should be designed with an awareness of cache coherence if speed is a primary consideration.

### 2.2 A Simple Genome Index

FastGA takes the approach of finding the adaptamer seeds between two genomes, obviating the need for selecting a *k*-mer size as it adaptively uses longer substrings in unique regions and shorter substrings in repetitive regions. In the original adaptamer implementation [26] a suffix tree of the second genome was built and then the first genome was scanned, finding the adaptamer hits at each position using the suffix tree. As discussed in Section 2.1 this is cache incoherent. Using a cache coherent approach, FastGA, effectively builds suffix trees of each genome and then in a novel merge algorithm finds the adaptamers of the suffixes in the first tree in the second. One of the advantages of this approach is that each suffix tree can be built independently for a genome, and then used in multiple comparisons with other genomes. But there is the problem of how to build a space-efficient suffix tree in a cache coherent manner. Also note that both the forward and complement strand of the genome must be represented.

Rather than build a suffix tree, first observe that almost all adaptamers for a multi-gigabyte genome will not be longer than than some reasonable choice of *K*, say 40, and for longer adaptamers their first *K* bases already constitute a very specific match into the second genome. So we simply build a sorted list of all the *K*-mers in the genome and its complement that occur fewer than *τ* times (user specifiable) and for each *K*-mer record the positions at which it occurs and its orientation, i.e. from the forward (+) or reverse strand (-), in the sign bit of the position. This is readily accomplished with a cache coherent MSD radix sort of (*K*-mer, signed-position) pairs as suggested earlier. As we will subsequently show, it is easy to have the MSD radix sort further return the *longest common prefix (lcp)* of a given *K*-mer and its predecessor in the final *K*-mer sorted array. Together the sorted *K*-mer list and lcp array provide a simple *genome index* that is in effect a suffix array that has been truncated at depth *K*. Given two such indices, we can find all the 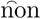-repetitive adaptamers of a suffix in the first index with respect to the second, where the adaptamer length is capped at *K*. Almost all genomes encountered are less than 100Gbp in size, and given this limit, a *K* = 40-mer is generally more than long enough to specifically correlate two regions of interest. A constant larger than 40 can be specified at compilation, but we have yet to encounter an example where empirically a larger choice of *K* is beneficial.

For a genome of length *N* , roughly 26*N* bytes are required for an index in which every 40-mer in both the genome and its complement are represented, a rather large data structure. To ameliorate this we can keep only those *K*-mers whose first *s* bases are a closed (*s, m*) syncmer for some choice of *s* and *m < s*. In practice we use (*s, m*) = (12, 8) effectively reducing the number of *K*-mers by a bit more than half. Note carefully that the complement of a syncmer is also a syncmer so in matching regions it is likely that both the forward and reverse adaptamers are synchronized implying only a slight loss in sensitivity. It further means that adaptamers of length less than *m* are lost, but again for *s* = 12 there are relatively few of these and they are not generally correlated with true matches.

Within FastGA *K* = 40, *s* = 12, and *m* = 8 are compile time parameters that could be retuned by a sophisticated user, e.g. a pair of very large, highly repetitive genomes, or lots of small, say bacterial, genomes, for a bit more performance. These selected values result in indices that occupy about 11GB per 1Gbp of genome and take about 15 seconds to construct on our laptop^1^ per 1Gbp of genome, using 8 threads. A final note is that merging two indices is a linear scan of both, as seen in Section 3, implying that the indices can be paged into memory as they are swept linearly thus requiring a very small memory footprint.

### 2.3 Most Significant Digit (MSD) Radix Sorting

In the last decade a number of researchers have discovered the power of most-significant-digit (MSD) radix sorting as a linear time, in-place, cache coherent sort of very large numbers of fixed size records [27, 28, 29]. Consider an array *A*[0..*N*− 1] of *N* elements each consisting of *K* bytes. View each element as a string of length *K* over an alphabet of size Σ = 256. An MSD radix sort, starts by sorting the elements in place on the first byte/symbol, partitioning the array into Σ sub-arrays where all the elements in each sub-array have the same first byte. Recursively, each of these sub-arrays is then sorted on the second byte, further partitioning each into Σ parts, which in turn are each sorted on the third byte and so on until depth *K* is reached at which point the elements are sorted. In a very simple form the in-place sort is as follows:

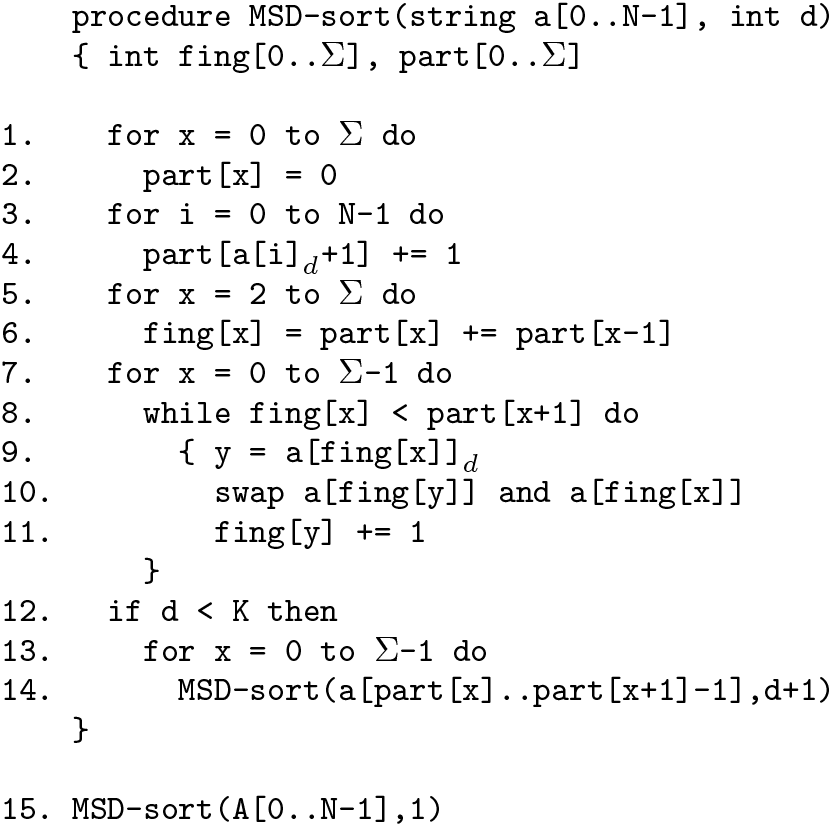

The first thing to observe is that the algorithm is cache-coherent as in any of its for-loops at most Σ + 1 regions of memory are being swept at any time. The number of steps is *O*(*NK*) but one must be careful as *O*(*K*) memory is being moved in step 10. for a complexity of *O*(*NK*^2^). However, for any fixed data size the sort is *O*(*N*). Further observe that in effect any data can be sorted. For example, a 64-bit integer can be viewed as an 8 byte string over an alphabet of size 256. As another example, we compress 40-mer DNA strings into 10 bytes using 2-bits per base, and sort (40-mer, contig, position) records to produce the genome indices of the previous subsection. Typically, the contig number occupies 2 bytes and the position within the contig 4, so that each record is 16 bytes. The array of such records is then MSD sorted on the first 10 bytes encoding the 40-mer. Another advantage of an MSD sort is that carefully memory mapping the array to be sorted gives one an effective external sort that can work on an arbitrarily large file of data.

The simple form above is usually refined in several ways. First, instead of just swapping two records, the implied permutation of the fingers is determined up to some maximum length, and then the records along the chain are moved in a cyclical fashion roughly halving the amount of data movement. Also when at depth *d* only the last *K* − (*d*− 1) bytes of each record need to be moved as the first *d*− 1 bytes of every record in the current sub-array are identical, again roughly halving the number of bytes moved. The second refinement, is that when a partition becomes small enough, a more efficient sort for this size is employed. For example in RADULS [27], introspective sort is used for *n* ≤ 384, shell sort for *n* ≤ 100, and insertion sort for *n* ≤ 32. Lastly, it is generally desirable to distribute the sort over multiple threads. This is easy to do once the first digit has resulted in Σ segments that can be sorted on the remaining digits independently. But the top level sort is not so easily distributed and fairly complex solutions that achieve good parallelism can be found in [27] and [29].

Our MSD sort departs from previous work in the following ways where the design assumes that a modest number *T* of threads, e.g. 4 to 16, are available. First, in the context of FastGA, we found it readily possible to know the sizes of each top-level partition and the amount a given “load thread” will contribute to each, so that each of *T* load threads, handling balanced amounts of the data, can construct their array records in locations so that the top level sort is already performed when the loading is complete. We then very simply divide the 256 top level partitions as evenly as possible across the *T* threads and let each thread proceed independently. We thus completely avoid the problem of threading the top level sort and postulate that this circumvention is possible for most applications.

Second, observe that while the partitioning is usually into Σ = 256 parts for small *d*, as the depth increases, there are an increasing number of empty partition intervals especially when *d* becomes larger than log_Σ_ *N* . Indeed, the number of non-empty partition intervals trends towards 1. So during the partitioning step (Lines 1-6 above) if we observe that only 1 partition interval is non-empty, we non-recursively increase *d* by 1 and repeat the partition step, as the *d*^*th*^ digit is equal for all entries. Thus, if a block of identical array elements occurs (e.g. ∼50 identical k-mers in a 50× shotgun data set), then this modification results in simply verifying that all the elements are equal and hence sorted in linear time with no data movement. In all other cases, we detect the *p* partition intervals that are non-empty and only iterate over their *p* digit values as opposed to all Σ. As one approaches small problems, *p* tends to be very small, 2 or 3, so that with this modification we find that our MSD sort is competitive without resorting to the special terminal sorts of previous approaches.

A third, subtle modifiaction is to detect when an entry is already in its correct partition so that it is not moved. This simple test further reduces data movement by roughly 10% overall. But note that for the deep cases, where only 2 or 3 partition intervals are non-empty, it results in 50% and 33% less data movement, respectively. This also further explains why a distinct terminal sort is not needed in our realization of the sort.

Our final refinement, especially important to the adaptive seed merge of Section 3, is to record the longest common prefix (lcp) of each element in the sorted array with its immediate predecessor. Simply, when sorting a sub-array on the *d*^*th*^ digit, the first element of each partition interval, except for the first element of the sub-array, has an lcp of *d* with its predecessor. If one is sorting 2-bit packed DNA strings a byte at a time, then the lcp of the first element of partition *p* with its predecessor is 4*d* + 3 − ⌊ log_4_ *p*∧*q* ⌋ assuming its predecessor is in partition *q < p* (remember, partition intervals can be empty), where *p*∧*q* is the bitwise exclusive-or operator. FastGA radix sorts on bytes (i.e. Σ = 256) and as the top level partition is recorded separately, we can conveniently stash the lcp for a genome index 40-mer in the first byte of each element.

### 2.4 Finding Seed Chains and Alignments

We defer the algorithm for merging two genome indices (with lcps) to Section 3.1. For the moment, in order to complete the overview of FastGA’s operation, we assert that the result of this merge is a set of pairs of positions in the two genomes and a length, (*i, j, t*), such that the string of length *t* starting at *i* is an adaptamer that matches the string of the same length starting at *j* in the second genome. In reality, each position say *i* is coded as a contig number *c*_*i*_ and then a position *p*_*i*_ within the contig. What we are seeking, in the dynamic programming matrix of contig *c*_*i*_ versus contig *c*_*j*_, is a *chain* of adaptamer seed hits that lie in a narrow diagonal band of width say *D* that are fairly tightly spaced within that band, say no two seeds are further apart than *A* anti-diagonals.

To find these chains efficiently, we create a list of records (*c*_*i*_, *c*_*j*_, ⌊(*p*_*i*_−*p*_*j*_)*/D⌋, p*_*i*_+*p*_*j*_, (*p*_*i*_−*p*_*j*_) *mod D, t*) for each adaptamer seed hit ((*c*_*i*_, *p*_*i*_), (*c*_*j*_, *p*_*j*_), *t*). In terms of the dynamic programming matrix of contig *c*_*i*_ versus contig *c*_*j*_, observe that the points (*p*_*i*_, *p*_*j*_) for which *l*(*p*_*i*_ − *p*_*j*_)*/Dj* = *b* for a fixed value of *b* ∈ [−|*c*_*j*_|*/D*, |*c*_*i*_|*/D*] are all in a diagonal band of width *D. p*_*i*_ + *p*_*j*_ is the anti-diagonal for the pair, and the exact diagonal for the pair can be reconstructed because the remainder of *p*_*i*_ − *p*_*j*_ modulo *D* is kept in the second to last entry. To recap, given a list entry (*c*_*i*_, *c*_*j*_, *b, a, r, t*) both *p*_*i*_ = (*a* + (*Db* + *r*))*/*2 and *p*_*j*_ = (*a* − (*Db* + *r*))*/*2 are indirectly encoded therein.

We then MSD sort this list of sextuplets on their first 4 components in lexicographical order. Thereafter, all the seed hits between a pair of contigs *c*_*i*_ and *c*_*j*_ are grouped together in the sorted list. Moreover, the seeds in each diagonal band are consecutive and in order of their anti-diagonal within the band. Thus chains of seed hits in a given diagonal band *b* that are not separated by more than *A* anti-diagonals are easily found in a linear scan of the sorted list.

However, we seek chains that can occur in *any* band of width *D*. To do so, the seeds in each pair of adjacent bands, *b* and *b* + 1, are merged in order of anti-diagonal and chains are sought in the merged output. This is still linear in time and captures all chains in a band of width *D*, but does also admit chains of width up to 2*D*. One could take further effort to only report chains of width *D* or less, but keeping in mind that the goal is to find pairs of regions that align, these extra wide chains only adversely affect search time (slightly) but not the sensitivity of FastGA. As chains are found in the linear sweep of the sorted hit list, FastGA tries to find a proper local alignment that passes through the *tube* of each chain. The tube is the rectangular region of the d.p. matrix that is bounded by the lowest and highest diagonals, *d*_*low*_ and *d*_*high*_, amongst the seeds in the chain, and the lowest and highest anti-diagonals, *a*_*low*_ and *a*_*high*_, where both ends of the seed hits are considered. This tube is easily derived as each chain is discovered.

FastGA uses the very rapid wave-based algorithm for finding local alignments that we reported previously in 2014 ([1]), so here we only describe the interface to it and will refer to it as the LA-finder. Given an anti-diagonal and an upper and lower diagonal bound, LA-finder finds the best local alignment that passes through the anti-diagonal between the diagonal bounds. It reports the end-points of this alignment, the number of differences within it, and a trace-point representation of the entire alignment (see Section 4). LA-finder always returns an alignment, which can have length 0 if there is no significant similarity in the region.

For each tube of a chain, LA-finder is first called on the anti-diagonal *a*_*low*_ + 2*D*, or (*a*_*low*_ + *a*_*high*_)*/*2 if *a*_*low*_ + *D* ≥ *a*_*high*_. If the alignment returned is sufficiently long and sufficiently similar (both parameters that can be set by the user) then the alignment is collected for possible output. Suppose the anti-diagonal at the far end of the alignment returned by LA-finder is *a*_*top*_. If *a*_*top*_ ≥ *a*_*high*_ then the tube is considered processed. Otherwise *a*_*low*_ is set to *a*_*top*_ and the now truncated tube is searched again. This continues until the entire tube has been searched with calls to LA-finder.

All the alignments for a given pair of contigs are collected and redundant alignments are removed. Exactly the same alignment or a prefix or suffix of a longer alignment can be found when disjoint chain tubes for the same alignment are found. Only one instance of the full alignment is kept. The *bounding box* of an alignment is the smallest rectangle that contains all the points on the alignment path. We further consider an alignment to be redundant if it is entirely within the bounding box of another alignment. For example, satellite repeats create waves of alignments that all fit in the bounding box of the alignment closest to the main diagonal. Finally, the resulting set of non-redundant alignments are output in order of their start coordinate in the first genome.

## 3 Algorithms

### 3.1 Finding Adaptamers

This subsection presents perhaps the most interesting algorithm within FastGA, where two genome indices, say 𝒢 and ℋ , of genomes *G* and *H*, are “merged” to find all the adaptamers of *G* and their matching positions in *H*. Previous work has shown that given a suffix array and its associated lcp array, one can build three “enhancing” arrays in linear time, such that one can then perform any operation possible on a suffix *tree* ([30]). This result transfers naturally to truncated suffix arrays. When viewed as suffix trees, our problem is in essence to find the intersection of the two suffix trees: a string is an adaptamer of *G* if it is spelled on a path to a vertex in the intersection that has an edge which is in the suffix tree of *G* but not *H*. This can be accomplished in time proportional to the number of edges in the uncompressed suffix trie of the intersection. However, the genome indices are very large objects, so building additional arrays to facilitate tree operations is prohibitive. So instead we developed an algorithm with the same time complexity that performs the task directly on 𝒢 and ℋ with only *O*(*K*) auxiliary storage, where *K* (= 40 in our implementation) is the length of the *k*-mers in both indices.

The algorithm processes each *K*-mer in the first index in sequence, which by the design of the indices are in sorted order. Formally, let 𝒢 [*i*].*kmer* be the i’th *K*-mer in the index, 𝒢 [*i*].*lcp* = *lcp*( 𝒢 [*i* − 1].*kmer*, 𝒢 [*i*].*kmer*) } be the lcp of the i’th *k*-mer with its predecessor, and let 𝒢 [*i*].*loc* = {*j* : *G*[*j, j* + *K*] = [*i*].*kmer* be the set of positions at which the i’th *K*-mer occurs in the source genome *G*. The problem then is for each *K*-mer in the first index, say *α* = 𝒢 [*i*].*kmer*, to find the range [*fst*_*i*_, *lst*_*i*_), *fst*_*i*_ *< lst*_*i*_ in the second index and length *L*_*i*_ such that *lcp*(*α*, ℋ [*j*].*kmer*) = *L*_*i*_ for all *j* ∈ [*fst*_*i*_, *lst*_*i*_) and *lcp*(*α, ℋ* [*j*].*kmer*) *< L*_*i*_ for all other *j*. Clearly, the adaptamer at each position *p* in 𝒢 [*i*].*loc* is the string *α*[0, *L*_*i*_] = *G*[*p, p* + *L*_*i*_] and each of these adaptamer instances matches at the locations *q* in 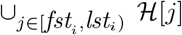. *loc* in the second genome. These pairs (*p, q*) of positions are appended to the match list for subsequent sorting as in the outline of Section 2.1 as long as the adaptamers are not repetitive, i.e. 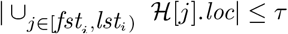

To find the range and length in ℋ, we need to merge the two indices in sorted order of *K*-mer. That is, for *K*-mer *α* at index *i* in *G*, we find the index *cur*_*i*_ in ℋ such that ℋ [*cur*_*i*_ − 1].*kmer < α* ≤ *H*[*cur*_*i*_].*kmer*. Immediately, observe that a longest prefix match to *α* in ℋ must occur at either *cur*_*i*_ − 1 or *cur*_*i*_, i.e. *L*_*i*_ = *max*(*lcp*(*α*, ℋ [*cur*_*i*_ − 1].*kmer*), *lcp*(*α*, ℋ [*cur*_*i*_].*kmer*)). Furthermore, *cur*_*i*_ must be in the interval [*fst*_*i*_, *lst*_*i*_]. Letting *c* be any index in [*fst*_*i*_, *lst*_*i*_), we can use the *lcp* information to quickly determine *fst*_*i*_ = *max*_*j*≤*c*_ ℋ [*j*].*lcp < L*_*i*_ and *lst*_*i*_ = *min*_*j*>*c*_ ℋ [*j*].*lcp < L*_*i*_. So intuitively if we can merge the two indices in order of *K*-mers we can use the *lcp*’s to find the desired intervals and adaptamer lengths. The task remaining is to do so as efficiently as possible.

Inductively, we seek to find *cur*_*i*_, [*fst*_*i*+1_, *lst*_*i*+1_), and *L*_*i*+1_ given these values for index *i* in 𝒢. To make the computation a bit more efficient, we will also maintain a *K* element vector *wall*_*i*_ such that for all *k < L*_*i*_, *wall*[*k*] = *min*_*j*_ *lcp*(*α*, ℋ [*j*].*kmer*) ≥ *k*. Figure 1 sketches the relationship of these variables in the second index ℋ . Observe that if *cur* = *lst* then ℋ [*cur*].*lcp < L* and if *cur < lst* then either *L* = *K* or ℋ [*cur*].*kmer*[*L* + 1] > *α*[*L* + 1]. The algorithm in Figure 2 below gives the logic for updating the invariant variables *fst, lst, L, cur* and *wall* from those for the *i*^*th*^ entry of 𝒢 to its *i* + 1^*st*^ entry. This logic is explained in the proof of correctness that now follows.

**Figure 1:**
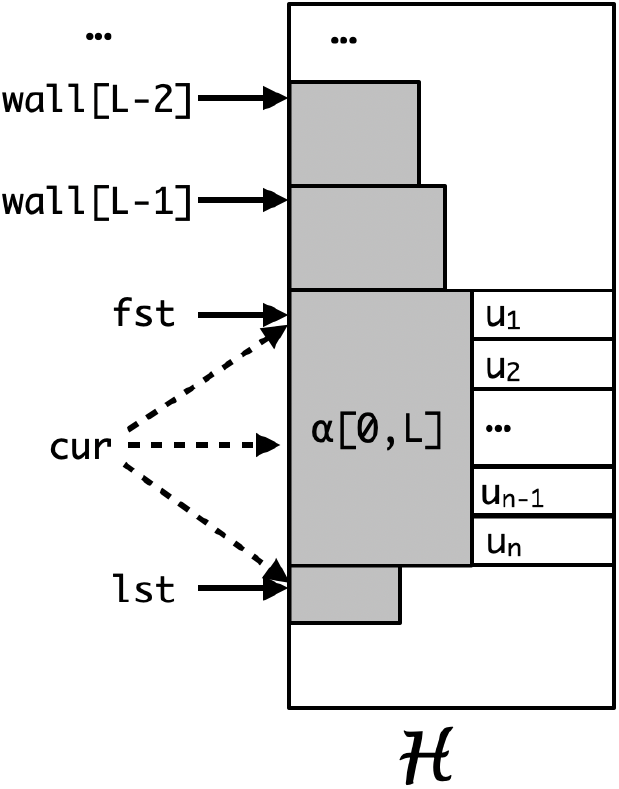
A sketch of the relationship of the variables *fst, cur, lst, L*, and *wall*. Assuming *α* is the current *K*-mer in the first index, then in the second index ℋ [*j*].*kmer*[0, *L*] = *α*[0, *L*] for all *j* ∈ [*fst, lst*). All the shaded areas have identical characters. The *lcp* of ℋ [*lst*].*kmer* with *α* is always strictly less than *L. cur* ∈ [*fst, lst*] points at the first element whose value is not less than *α* and can only equal *α* when *L* = *K, fst* = *cur* ,and *lst* = *cur* + 1. If *L < K* then the *L* + 1^*st*^ symbols in the adaptamer interval are strictly increasing, i.e., *u*_1_ ≤ *u*_2_ ≤ … ≤ *u*_*n*_ and if *cur < lst* then *α*[*L* + 1] *<* ℋ [*cur*].*kmer*[*L* + 1] and if *cur* > *fst* then ℋ [*cur* − 1].*kmer*[*L* + 1] *< α*[*L* + 1]. The *wall* values for *l < L* give the smallest index for which the *K*-mer has an *lcp* of *l* with *α*. |*wall*[*l*] − *wall*[*l* − 1]| can be zero.

**Figure 2:**
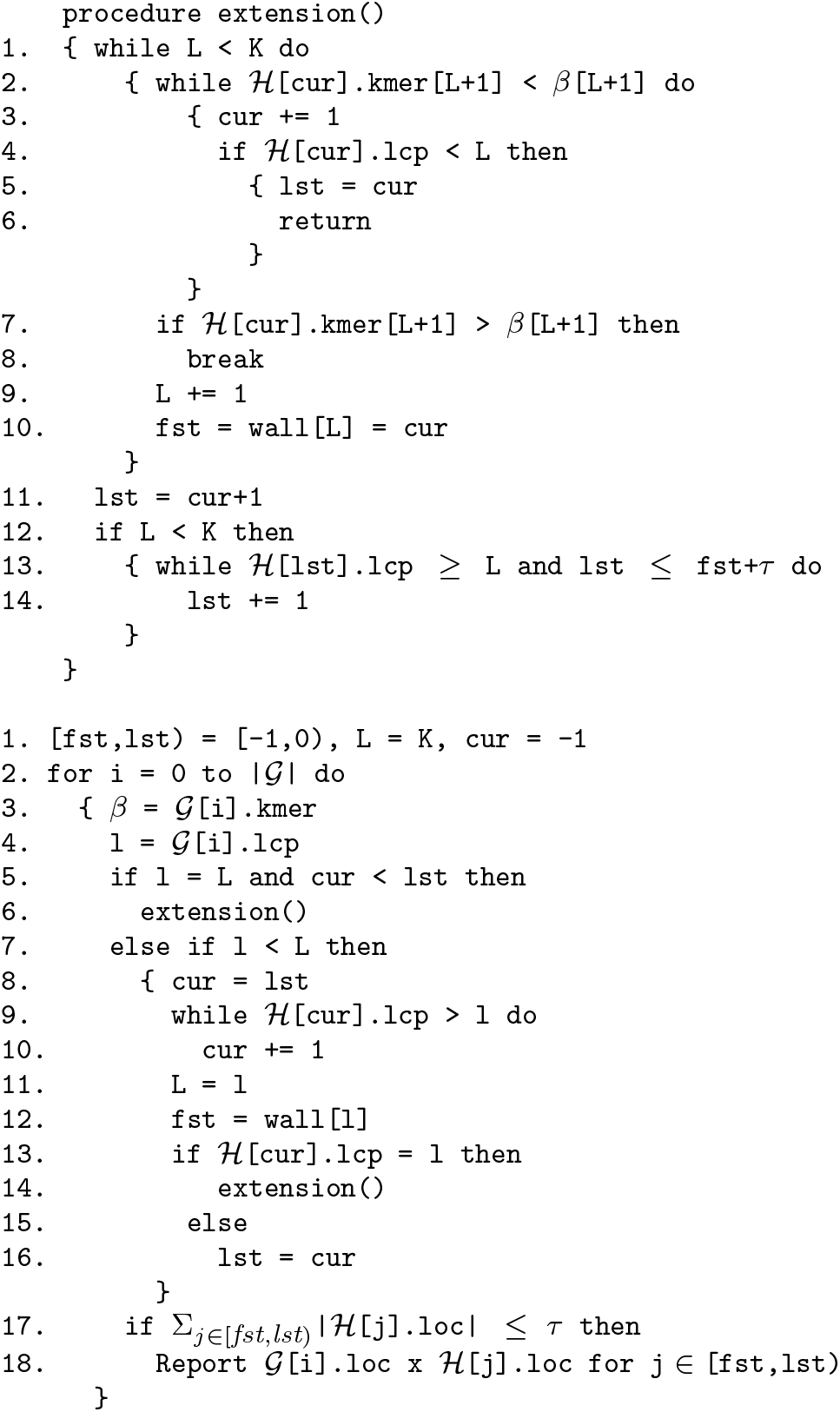
An algorithm for finding all the non-repetitive adaptamers of genome G in genome H given their indices 𝒢 and ℋ.

#### Theorem 1

The algorithm in Figure 2 finds all non-repetitive adaptamer matches in *G* to positions in *H* in a sweep of their indices 𝒢 and ℋ.

**Proof**:

To start the induction imagine that both indices begin with a special *K*-mer that consists entirely of a special symbol ζ that is not in the alphabet of the indices and is smaller than any symbol in said alphabet. Then clearly, the correct values for this −1^*st*^ entry of 𝒢 is [*fst, lst*) = [− 1, 0), *L* = *K* and *cur* = −1 (Line 1).

The algorithm then proceeds to find the correct values for each index *i* in 𝒢 in order (Line 1) in the body of the for loop incrementing *i* (Lines 3 to 16) and further outputs the adaptamer matches for the locations of *i* if not repetitive (Lines 17-18).

So consider then that *fst, lst, L, cur* and *wall* have the correct values for index *i* − 1 as the loop of lines 3-18 is entered with index *i*. To ease notation let *α* = 𝒢 [*i*− 1].*kmer, β* = 𝒢 [*i* + 1].*kmer*, and *l* = 𝒢 [*i* + 1].*lcp*. If *l* > *L* then the answer for both *i* and *i*+1 are the same as *β*[0, *L*+1] = *α*[0, *L*+1] and a match longer than *L* to *α* is already known to not be possible. If *l* = *L* and *cur* = *lst* then *β*[0, *L*] = *α*[0, *L*] *<* [*lst*].*kmer*[0, *L*] so again a match longer than *L* to *β* is not possible, so the variables remain unchanged. On the other hand if *l* = *L* and *cur < lst* [Lines 6-7] then a longer match of *β* to ℋ [*j*].*kmer* for some *j* ≥ *cur* is possible and it is sought in the procedure extension. Lastly, if *l < L* [Line 9] then *β*[0, *L*] > *α*[0, *L*] implying *cur* ≥ *lst* (Line 8). Moreover as long as ℋ [*cur*].*lcp* > *l, β*[0, *l* + 1] > *α*[0, *l* + 1] = ℋ [*cur*].*kmer*[0, *l* + 1] and so *cur* should be advanced (Lines 9-10). If after advance *cur*, ℋ [*cur*].*lcp < l* then *beta*[0, *l*] *< H*[*cur*].*kmer*[0, *l*] implying *β <* ℋ [*cur*].*kmer* and *β*[0, *l*] is the longest match of *β* to the preceding values of *cur*. So *cur* = *cur*_*i*_ = *lst*_*i*_ and *fst*_*i*_ = *wall*[*l*] and *L*_*i*_ = *l*, so lines 11, 12, 15 and 16 are correct. Note in this case, the adaptamer range is a superset of the range for *i* − 1 with a smaller adaptamer length. On the other hand, if after advance *cur*, ℋ [*cur*].*lcp* = *l*, then *β*[0, *l*] = ℋ [*cur*].*kmer*[0, *L*] so a longer match is possible and sought by entering procedure `extension`.

Upon entry to `extension`, ℋ [*cur* − 1].*kmer < β*, and *β*[0, *L*] = ℋ [*cur*].*kmer*[0, *L*] = [*fst*].*kmer*[0, *L*] > ℋ [*fst* − 1].*kmer*[0, *L*]. The first goal is to advance *cur* until ℋ [*cur*].*kmer* ≥ *β*. If *L* = *K* (Line 1) then we are done as it implies *β* = ℋ [*cur*].*kmer*. Otherwise if ℋ [*cur*].*kmer*[*L* + 1] *< β*[*L* + 1] then ℋ [*cur*].*kmer < β* so *cur* should be advanced to the next index (Lines 2-3). If the *lcp* at this next element is less than *L*, then [*cur*].*kmer*[0, *L*] > [*cur* − 1].*kmer*[0, *L*] implying [*cur*].*kmer* > *β* and moreover that the adaptamer length of *β* is *L* and *lst* = *cur* (Lines 4-6). The routine returns in this instance. Otherwise it continues the loop of Lines 2-6 with this new value of *cur*. If this loop exits then ℋ [*cur*].*kmer*[*L* + 1] ≥ *β*[*L* + 1]. If ℋ [*cur*].*kmer*[*L* + 1] > *β*[*L* + 1] then we have found the new value of *cur* as ℋ [*cur*].*kmer* > *β* and the extension loop of Lines 1-10 is exited. Otherwise the *L* + 1^*st*^ symbols are equal, so *L* is increased by 1 and *wall*[*L*] is updated and is potentially the new value of *fst* unless further extensions occur in the loop (Lines 9-10).

So when (and if) the extension loop of Lines 1-10 exits ℋ [*cur*].*kmer* ≥ *β, L* = *lcp*(*β*, [*cur*].*kmer*) is the adaptamer length for *β*. In this instance *lst* > *cur* as *cur* is within the range of indices in ℋ that match the adaptamer. If *L* = *K* then this range is just the index *cur* and *lst* = *cur* + 1 (Lines 11-12). Otherwise, we must search forward from *cur* to find the last index in ℋ that matches the adaptamer prefix of length *L*. As before this is simply a matter of finding the first index for which its *lcp* is less than *L* (Lines 13-14). There is in the worst case no strong bound on how many indices would need to be searched, so pragmatically we cap *lst* at *fst* + *τ* + 1 as this implies that the adaptamer is repetitive and so its pairs will not be reported in Lines 17-18 of the main loop. Revisiting the early parts of the proof one can see that this slight change in the inductive hypothesis, i.e. either *lst* has its desired value or is *fst* + *τ* + 1 if it is greater, does not affect the correctness of the algorithm. ▪

Considering this algorithm in terms of intersecting two suffix tries, it takes time proportional to the number of edges in the intersection, save for vertices near the root where *O*(*τ*) time might be taken as opposed to *O*(1) time. Theoretically we could just consider *τ* a small constant. But to get an idea of how much work is done in a typical merge, we captured statistics for our running example of comparing two bats where the genomes are of size 2.23Gbp (*G*) and 2.56Gbp (*H*). 𝒢 has *N* = 1.603 billion K-mers that begin with (12, 8) syncmers. The intersection of this index with ℋ had only 2.646 billion edges indicating that most consecutive adaptamers differ in one or two characters at their end, that is, the intersection is *O*(*N*) in expectation (and not *O*(*KN*), the worst case). Moreover, the algorithm only makes 4.356 billion character comparisons, or less than 3 per adaptamer found. Finally, line 14 was executed only .807 billion times reflecting that in most cases the (*fst, lst*) interval is typically of size 1. While only a single data point, this is typical in our experience, and exemplifies why it is much more efficient to merge indices rather than to do single entry lookups.

The fact that an adaptamer in *G* is not necessarily an adaptamer in *H* gave us some concern. One can make matters symmetric by simply computing the union of the seed hits for the adaptamers of both genomes and FastGA will do this when the -S option is set. We achieve this by simply running the merge of 𝒢 versus ℋ and then of ℋ versus 𝒢 , and taking the union of the seed locations found. We anecdotally observe that the difference is primarily that, with the union, FastGA finds additional short repetitive alignments in *H*. Thus for genome alignment where repetitive matches not flanked by homologous sequence are not of interest, we advise that the difference is not important enough to be worth the additional compute time. The one application where we would recommend using the -S option is for *ab initio* repeat finding and modeling.

### 3.2 Storing and Delivering Alignments

#### 3.2.1 Trace Points

FastGA typically finds anywhere from several hundred thousand to tens of millions of alignments depending on genome size, evolutionary distance, and repetitiveness. Each alignment aligns an interval *A*[*ab, ae*] of the first genome against an interval *B*[*bb, be*] of the second. To encode the alignment there are two obvious approaches: (1) just record the 4 interval endpoints and recompute the details of the alignment when needed, or (2) record the details by noting where exactly each inserted and deleted base occurs so that the alignment can be delivered in linear time, e.g. via a CIGAR string [2]. For this discussion assume *n* = *ae* − *ab* and the sequence dissimilarity is *ϵ* = *d/n*, so that *be* − *bb* ≤ *n*(1 + *ϵ*) = *o*(*n*) and we use *d* or *ϵ* interchangeably in characterizing complexities. So approach (1) requires *o*(1) space to store the alignment but *O*(*nd*) worst-case, *O*(*n* + *d*^2^) expected time to deliver it when needed using the Myers-Ukkonen wave algorithm [31, 32]. In contrast, approach (2) requires *o*(*d*) space for storage and only *O*(*n*) to deliver the alignment. Note in both cases the work-space product is *O*(*nd*).

The idea of *trace points* delivers a continuum of space-time tradeoffs between the two methods that is determined by a *trace point spacing* parameterδ. Consider the sequence *A* to be partitioned into a series of panels – *A*[0,δ], *A*[*δ*, 2*δ*], *A*[2*δ*, 3*δ*], … each of exactlyδ symbols. Then *A*[*ab, ae*] is the concatenation of the panels *A*[*ab, x*_1_*δ*], *A*[*x*_1_*δ, x*_2_*δ*], … *A*[*x*_*t*_*δ, ae*] where *x*_1_ = ⌊*ab/δ* + 1⌋ , *x*_*i*_ = *x*_*i*−1_ + 1, and *x*_*t*_ = ⌊ (*ae* 1)*/δ*⌋ . Obviously, the first and last panels may be shorter thanδ but the point is that the division of the interval [*ab, ae*] is implicit, one only needsδ to know the partitioning. Next, observe that in an alignment between *A*[*ab, ae*] and *B*[*bb, be*], the *B* interval *B*[*bb, be*] is itself partitioned into the same number of panels as the partition *A, B*[*y*_0_, *y*_1_], *B*[*y*_1_, *y*_2_], … *B*[*y*_*t*_, *y*_*t*+1_] where *y*_0_ = *bb, y*_*t*+1_ = *be*, and the overall alignment is the concatenation of the subalignments between *A*[*x*_*i*_*δ, x*_*i*+1_*δ*] versus *B*[*y*_*i*_, *y*_*i*+1_]. This *B* partitioning is not unique, as the splitting of any symbols inserted in *B* between the symbols 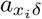 and 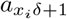 can be arbitrary. But all that is important is that such a division is possible and an instance easily determined (e.g. place all inserted *B* symbols in the left interval). Figure 3 illustrates the division. Immediately observe that the length of each *B* panel induced by the alignment, *b*_*i*_ = *y*_*i*+1_ − *y*_*i*_ for *i* = 0 to *t*, is not fixed but generally varies betweenδ(1 − ϵ) andδ(1 + ϵ). We call the *t* + 1-element array of *B*-panel lengths, *< b*_0_, *b*_1_, …, *b*_*t*_ > the *trace point array* for the alignment.

**Figure 3:**
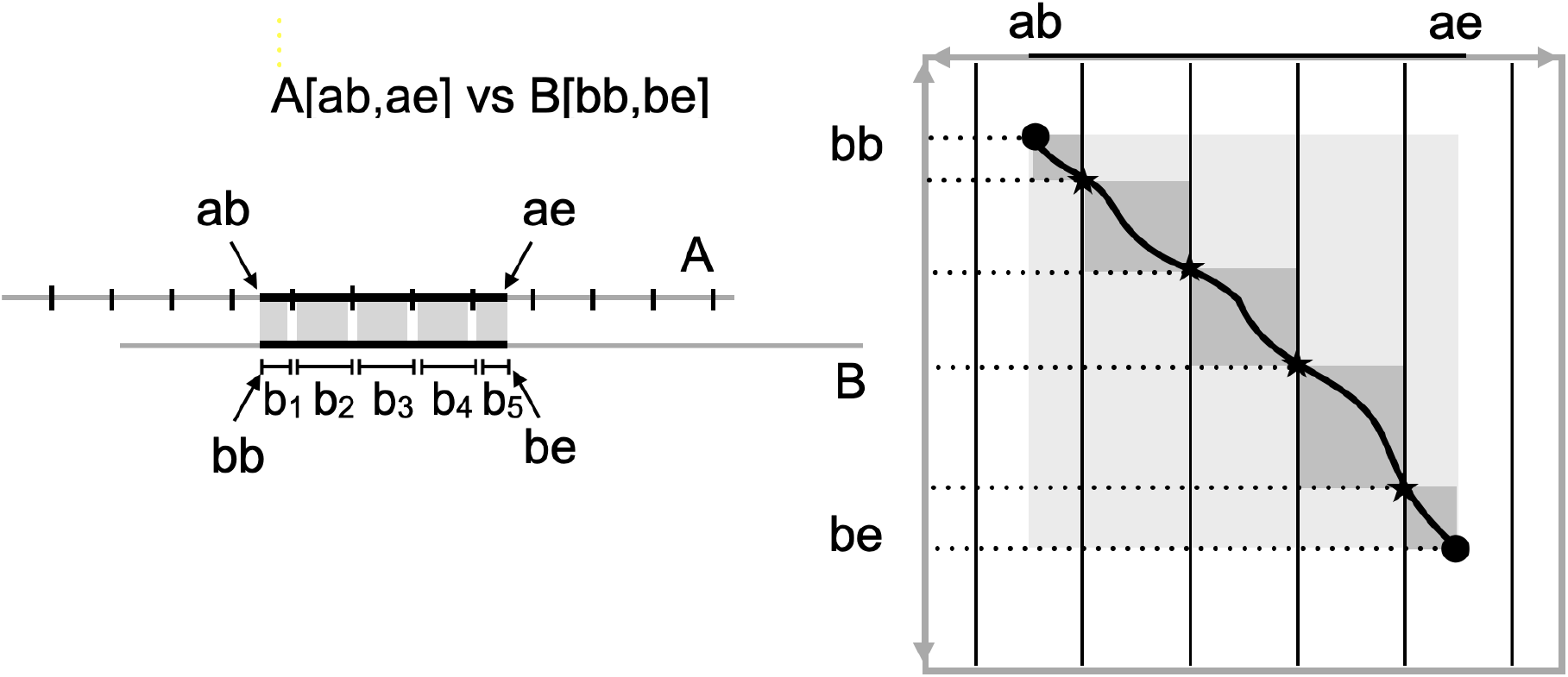
In the figure at left, *A* is divided evenly into panels of sizeδ save the last. For the alignment of *A*[*ab, ae*] with *B*[*bb, be*], the length of the portion of *B* aligned with each segment of *A* as partitioned by theδ-panels is the trace point array. Given this array, the 4 end points, andδ on can then determine each starred *trace-point* in the edit graph of *A* versus *B* which are vertices that the encoded alignment pass through. Given these points, delivering the path is simply a matter of solving each of *o*(*n/δ*),δ×*δ* sub-arrays shown in dark grey between the trace points.

The trace point array, along with the four endpoints, provides a record of an alignment with just *t* + 1 = *o*(*n/δ*) integers of magnitude *O*(*δ*), and we now show how to reconstruct the full alignment in *O*(*n* +δ*d*) time. Given the array one can easily in *O*(*n/δ*) time reconstruct the *y*_*i*_’s and *x*_*i*_’s above, so the problem remains to deliver the *t* + 1 sub-alignments for *A*[*x*_*i*_*δ, x*_*i*+1_*δ*] versus *B*[*y*_*i*_, *y*_*i*+1_] in order of *i*. For this we use the *O*((*n* +*m*)(*p* +1)) “skewed-wave” algorithm by Myers and co-authors from 1990 [33] where the parameter *p* = *d* − |*m* − *n*| reduces the effect of “skewed” problems where *n ≫ m* or vice versa. This is important here because theoretically there is no limit on the skew between a particular *b*_*i*_ andδ. Suppose *b*_*i*_ =δ + *x* for *x* > 0. Then the skew wave algorithm takes *O*(*δ* + (*δ* + *x*)(*d*_*i*_ − *x*)) where *d*_*i*_ is the number of differences for sub-problem *i* and thus Σ_*i*_*d*_*i*_ = *d*. At worst *d*_*i*_ = *b*_*i*_ under the simple unit-cost model, so *d*_*i*_ − *x* ≤ *b*_*i*_ − *x* =δ and thus (*δ* + *x*)(*d*_*i*_ − *x*) =δ*d*_*i*_ + *x*((*d*_*i*_ − *x*) −δ) ≤δ*d*_*i*_. So the time taken is *O*(*δ*(*d*_*i*_ + 1)). If *b*_*i*_ =δ − *x*, then the algorithm takes *O*(*δ*(*d*_*i*_ − (*δ* − *x*))) which is trivially *O*(*δd*_*i*_). So altogether the time taken to compute the *t* + 1 sub-alignments is *O*(Σ_*i*_*δ*(*d*_*i*_ + 1)) = *O*(*n* +δ*d*).

Note that the space-time product is *O*(*n/δ*)*O*(*n* +δ*d*) = *O*(*n*(*d* + *n/δ*)) which is a bit more than for the initial two approaches, but nonetheless we have achieved a continuum of space-time tradeoffs parameterized onδ. In particular, note that constructing an alignment from the tracepoints is *O*(*n*) for a fixed value ofδ and the space-time product *is O*(*nd*) when ϵ ≥ 1*/δ*. In FastGA and other applications we useδ = 100 as it allows the *b*_*i*_ to fit in a single byte. The astute reader will note that a trace point length could exceed 255 if there is a large insertion in *B* for a particular panel. One can handle this by using a coding scheme for integers that is adaptive to size so that a byte is used in expectation. But we have rather implemented our unit-cost wave algorithm so that it considers such a large gap a break in genome similarity as discussed earlier. As an example then, for an alignment covering 10,000bp we encode an alignment in 100 bytes regardless of ϵ, versus a CIGAR string of thousands of bytes for an 80% identity alignment. Moreover, in empirical trials we can convert our trace-point encoded ALN-files to PAF-format with CIGAR strings at a rate of roughly 100 million aligned bases per second on our laptop^1^ including the time taken to refine gaps as described next.

#### 3.2.2 Gap Refinement

While our adaptive wave algorithm is well-suited to discovering similar regions, it is true that the alignments it delivers do not properly align letters within even modest gaps, for example see Figure 4(a). An interesting empirical observation is that most of these undesirable placements would be rectified if one could deliver the optimal unit cost alignment with the *fewest gaps*, for example, see Figure 4(b). The connection to the use of an affine gap model for alignment is that solving this problem is equivalent to adding an infinitesimal gap start cost to the unit cost model^2^.

**Figure 4:**
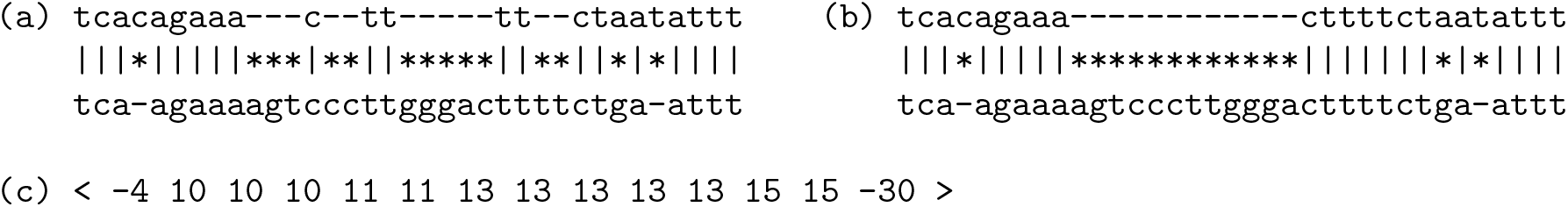
An example of gap refinement. At left, (a), an alignment produced under the unit-cost model. At right, (b) an alignment with the same unit cost but only one gap. In (c) we give the *trace* for the alignment (a), see the text for the explanation.

Therefore, prior to reporting an alignment’s path through the underlying dynamic programming (d.p.) matrix, we seek to modify the path so as to reduce the number of gaps. We encode the path delivered by the trace point algorithm described above as an *O*(*d*) array that indicates where to place dashes between the symbols of *A* and *B* so as to expose the aligned bases as illustrated in Figures 4 above. Specifically, we produce an *indel array < o*_1_, *o*_2_, ..*o*_*a*_ > of strictly positive and negative integers of non-decreasing magnitude where if *o*_*i*_ > 0 then one should insert a dash before symbol 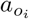 of the sequence *A*, and if *o*_*i*_ *<* 0 then one should insert a dash before symbol 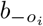 of the sequence *B*. Figure 4(c) gives an example of the indel array for the alignment of Figure 4(a). In this representation of the alignment, the regions we seek to improve are captured by maximal segments of the array *< o*_*x*_, *o*_*x*+1_, …*o*_*y*_ > such that (a) all the numbers have the same sign (i.e. all deletions or all insertions), (b) *y* − *x* ≥1, and (c) |*o*_*i*+1_ − *o*_*i*_| *< R* for all *i* ∈ [*x, y*), where *R* is a separation parameter that we set to 50. This is easy to do in a linear sweep of the array. Without loss of generality we consider the case where all numbers are positive. The region between the first and last insert consists of *D* = *y* − *x* + 2 diagonals and *L* = *o*_*y*_ − *o*_*x*_ + 1 columns of the d.p. matrix, which we consider in relative terms by considering the start of the insertion edge for *o*_*x*_ as the coordinate (0, 0). Figure 5 illustrates this *D* × *L* trapezoidal region for the positive run of the trace for Figure 4(a) and the revised path corresponding to the alignment in 4(b).

**Figure 5:**
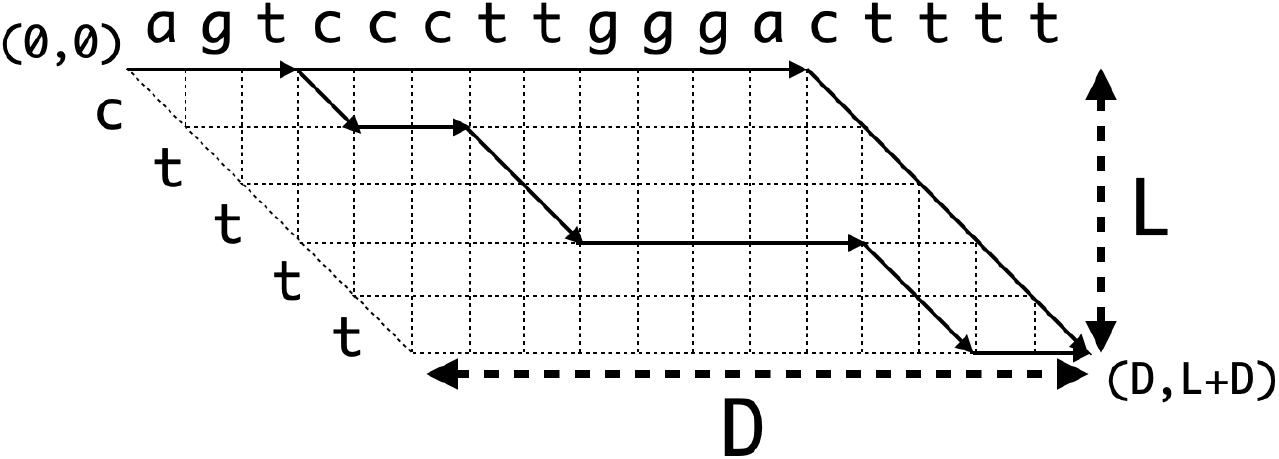
The trapezoidal region to be re-examined for the example of Figure 4 with the paths corresponding to alignments (a) and (b) drawn in black.

We seek the best alignment from (0,0) to (D,L+D) that does not involved deletions as we are only assessing whether the number of insertion gaps can be reduced. So our basic recurrence is the classic one of Gotoh but with the deletion or D-terms removed and a gap initiation cost of 1 (so a first insertion costs 2):

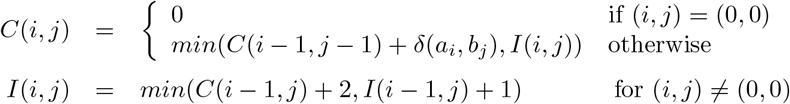

Rather than compute the recurrences above over the trapezoid, we use path compression [34] and a greedy wave approach to arrive at a novel and faster solution for this special problem. A simple lower bound on the cost of any path from a vertex (*i, j*) to the goal Φ = (*L, D* + *L*) is the number of insertions needed to get from its diagonal *i* − *j* to the goal diagonal *D*, i.e. *L*(*i, j*) = *D*− (*i* − *j*). Given such a lower bound and the *acyclic* d.p. graph, the cost of every edge *v* → *w* can be reduced by *L*(*v*) − *L*(*w*) without changing the set of optimal paths whose cost under the “compressed” problem is *C*^*1*^(Φ) = *C*(Φ) − *L*(Φ) = *C*(Φ) −*G* (see [34]). In our specific case, the given lower bound reduces the cost of insertion edges by 1, so that the recurrence for *I*-nodes becomes:

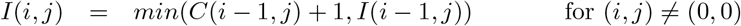

that is, gap starts cost 1 and gap continuations have 0 cost.

We find the optimal path under the compressed edge costs using a wave algorithm where the cost *k*-wave consists of the furthest *C*-vertices *F*_*k*_(*d*) reachable along each diagonal *d* ∈ [0, *D*] via a cost *k* path from the origin (0, 0). To determine the recurrence for *F*_*k*+1_(*d*) we need to consider all the ways one could reach the point on a path of cost 1 from a *k*-wave vertex. To wit, one can

a. start at *F*_*k*_(*e*) for *e < d*, take a cost 1 insertion to the *I*-node in diagonal *e* + 1, then take 0 cost insertion edges to diagonal *d*, transition to the *C*-node, and then follow 0-cost diagonal *C* edges as far as possible, or
b. start at *F*_*k*_(*d*), take a substitution edge of cost 1 forward along diagonal *d*, then follow 0-cost diagonal *C* edges as far as possible

This leads to the following recurrence:

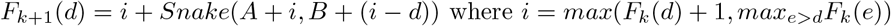

which in turn leads to the simple algorithm below where a single array F[0..D] keeps the furthest reaching points in each diagonal *d* by keep the *A* sequence position *i* where the *B* position is then implicitly *d* − *i*.

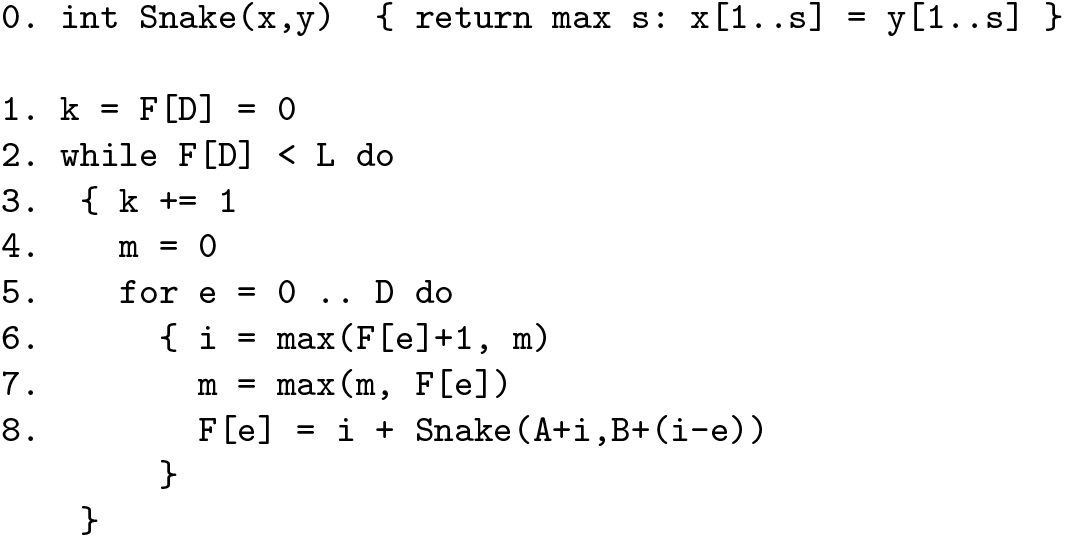

The algorithm stops with *k* = *C*^*′*^ (Φ) which is the number of gaps and mismatches in the alignment. If *G* is the number of gaps and *S* the number of mismatches in the best answer of score *k* = *G* + *S*, then the algorithm takes *O*((*G* + *S*)*D* + *L*) expected time. A bi-directed linear space construction of a best alignment achieving this score is *not* possible for the compressed version of the problem, so we simply maintain a vector for each successive wave and then trace back the optimal path involving the smallest number of gaps. Indeed, note that with the chosen scoring scheme, we will permit a single mismatch if it allows us to eliminate a gap. If we had chosen a gap start cost of *g*, then we would be permitting *g* mismatches in order to eliminate a gap. Although in theory gap costs can be associated with expected insertion distributions, in practice our experience is that choosing *g* = 1 behaves robustly to avoid obviously improvable gaps in alignments on visual inspection.

To get an idea of how much work is done processing a typical alignment file, we captured statistics for our comparison between two bats that delivers 1.63 billion base pairs of aligned sequence. There were 64.6 million gaps in the initial set of unit-cost alignments. 46.1 million of these occurred in 11.93 million inspected trapezoids for which we ended up removing 13.7 million gaps. The average product of (*G* + *S*)*D* was 88 over all the trapezoids, with the maximum for *D* being 121, the maximum for *G* + *S* being 133, and the largest product being 15, 125. 9.6 million of the problems had product less than 100, and 11.84*M* were less than 1000. When converting a trace-point alignment file of these alignments to CIGAR strings for a PAF file, only 6.7 seconds were spent on gap refinement which constituted roughly 15% of the total compute time for the task.

## 4 Software

### 4.1 The Framework

Using **FastGA** can be as simple as calling it with two (optionally gzip compressed) FASTA files containing genome sequences, where each entry is a scaffold with runs of Ns separating contigs. By default a PAF file encoding all the local alignments found between the two genomes is streamed to the standard output. The code is available at http://www.github.com/thegenemyers/FASTGA along with some example data so the user can try running it themselves.

Under the surface, a number of intermediate steps take place. First, the FASTA files are converted to *genome databases* with extension .`1gdb` that are a `ONEcode` [3] binary file and associated hidden file containing the ASCII DNA sequences in 2-bit compressed form. This allows FastGA to randomly access contigs without text parsing. Second, a *genome index* with extension `.gix` is then built for each genome that contains the truncated suffix array and associated lcp array (see Subsection 2.2). Third, FastGA records all the alignments it finds in a `ONEcode` binary file we refer to as an ALN-formatted file with extension .1aln that uses the space efficient trace point encoding of each alignment described earlier in subsection 4.1. Finally in linear time, this trace point representation is converted into the desired PAF output. Importantly, one has the option to keep the results in the ALN file, and then convert it to any of PAF, PSL, or other desired alignment format on demand. The diagram of Figure 6 summarizes and details the data flow just described.

**Figure 6:**
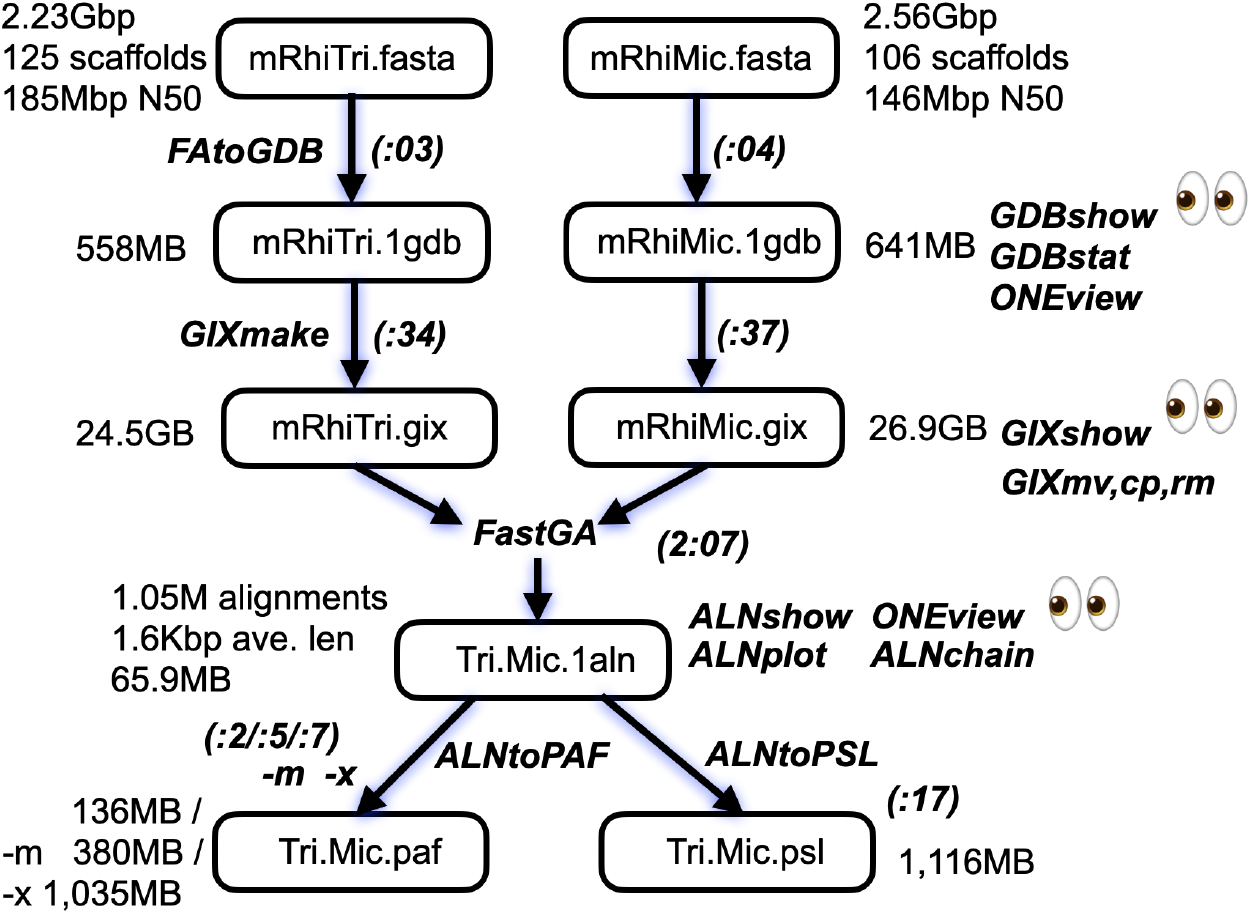
The data work flow for FastGA and associated utilities. Each box contains a data object and the arrows are labeled with program names that transform or produce them. Sample data sizes and parameters are placed to the side of each box and runtimes are placed in parenthesis along the program arrows. Utilities for viewing and or manipulating a data object are listed at the right with a pair of eyes next to those that provide views.

While the entire set of transformation processes can be fired off by simply calling FastGA, we provide routines to perform each step under direct control. In addition we provide utilities listed at right in Figure 6 that allow one to examine the intermediate GDB, GIX, and ALN files. An invocation of FastGA with the -k option or direct application of the sub-process routines, creates persistent GDB and GIX entities that can be reused, saving time if a given genome is to be compared repeatedly. The GDB and GIX items are actually each an ensemble, consisting of a proxy file and a number of hidden files. So we provide the utilities GIXmv, GIXcp, and GIXrm to manipulate these ensembles. Finally, we provide the utility GDBtoFA that inverts the process of converting a FASTA file into a GDB.

FastGA features the use of the `ONEcode` data encoding framework for both its’ GDB and ALN files that encode all the alignments found. As such FastGA also supports as input `ONEcode` sequence files that encode a genome, in addition to the usual FASTA format. So both FAtoGDB and GDBtoFA (despite their names) also recognize and support `ONEcode` SEQ files as well as FASTA. In addition, the GDB and ALN files can be viewed with the `ONEview` utility of the `ONEcode` framework.

### 4.2 ONEcode

The alignment records produced by FastGA are written into a `ONEcode` file [3]. `ONEcode` is a general data framework created by Myers and Durbin that is unpublished and in the open domain being freely available at http://www.github.com/thegenemyers/ONEcode. The three main ideas dominating the design of the framework are that (1) it should be easy to understand and use, (2) the core implementation should be succint without any package dependencies, and (3) despite (1) and (2) it should support space and time efficient binary representations fit for production scale data analysis. A key feature is that, in common with other bioinformatics formats such as SAM/BAM and VCF/BCF there are always interconvertible ASCII and binary versions of each ONEcode file.

In the ASCII version, data is stored as a sequence of strongly-typed lines (records) each of which begins with a single character (the 1-code) that determines the nature of the data that follows on the line, where individual items can be an integer, real, char, or list of any of these. Integers and reals can be up to 64-bit precision and only one list is permitted per line type. Every `ONEcode` file has a header section that (a) gives the series of commands, or provenance, that created it, (b) the schema of the data to follow, i.e. it is self-describing, and (c) information on the number and maximum sizes of the various types of data that follow, in support of aim (1) above. The ASCII encoding is normally only used for viewing by a human being or rapid prototyping at a small scale. For every ASCII file there is a corresponding binary encoding which (a) permits random access to any item via an indexing scheme, and (b) compresses every item in a manner specific to the datum’s type and on a per item basis.

In support of the framework, there is a C-library that reads and writes `ONEcode` data, where reading is transparent to whether the input is ASCII or binary and writing can be in either mode. It further supports multi-threaded reading and writing. There are also two utilities: `ONEview`, that allows extractions of arbitrary (intervals of) objects and also can convert between binary and ASCII file types, and `ONEstat`, that verifies the correctness of a `ONEcode` file and can display summary statistics about what is within.

In our experience, `ONEcode` binary files are as efficient if not more so than say gzip’d FASTA files or BAM files, while allowing direct indexing without the need of precomputing an index of a file (e.g. bgzip). Indeed, in the case of storing alignments we had originally been using a custom binary format for storing trace-point based alignment records, but when we developed an equivalent `ONEcode`schema, the resulting files were 25% smaller due to the built-in line compression. Manipulating and viewing these files is then immediately supported by the framework at production scale efficiency. In particular, the schema entails encoding each tracepoint sequence as a list of integers. `ONEcode` stores these as the first value and then the sequence of first-forward differences in the minimum number of bits possible, and then compresses these with a Huffman scheme trained on the condensed trace-point lists. New schemas for `ONEcode` file types can easily be constructed for new data models, and they can also be extended while maintaining backwards compatibility with existing ones. Indeed we have already used `ONEcode` files for diverse purposes including in the ancient DNA phylogeny placement package PathPhynder [35].

## 5 Experimental Results

In order to assess the performance of FastGA, we compared it to four other genome aligners: minimap2 ([19], version 2.28), LastZ ([16], version 1.04.22), wfmash ([22], version 0.21.0) and NUCmer ([18], version 4.0.0rc1). We first evaluated the performance of each method using simulated genomes to establish a controlled benchmark. We then applied the methods to real genomic data by aligning several mammalian genomes to the CHM13 human reference ([36]), assessing performance in a more biologically relevant context. Finally, to evaluate the generalisability of the methods across divergent species, we extended our analysis to representatives from six animal lineages across a range of genome sizes: one each from insects, fishes, birds, reptiles, mammals, and amphibia.

For real genomes, the soft-masked sequences were supplied to LastZ, allowing the tool to appropriately handle repetitive regions. We enabled alignment CIGAR-string output by adding the ‘-pafx’ option to FastGA, ‘-c’ option to minimap2 and ‘--format=PAF:wfmash’ option to LastZ. For LastZ and NUCmer, alignments were performed between individual pairs of chromosomes (or other assembly scaffold sequences) to accommodate input sequence size limitations, with each job executed using a single thread; while for FastGA, minimap2 and wfmash, the entire genomes were used as inputs and the alignments were performed using 16 threads. CPU time and memory usage for each job were recorded using the Linux command ‘/usr/bin/time -v’. Experiments were performed on a machine running the Ubuntu 22.04 Linux operating system, equipped with 376 GB of memory and two Intel(R) Xeon(R) Gold 6226R CPUs @ 2.90GHz with a total of 32 cores supporting up to 64 threads.

## 5.1 Simulated Genomes

We simulated a pair of genomes, designated 𝒜 andℬ , composed of 10 kb blocks, each starting with a region of similarity of varying length (from 100 bp to 5 kb) and divergence (from 1% to 65%) followed by random sequence, with the order of the blocks randomised in the genomes so that there are no long-range alignments across multiple blocks. With 100 replicates for set of similarity parameters (length and divergence) in total the “genomes” were each 84 Mb long. The sequence divergences were introduced by random single nucleotide substitutions (80%), insertions (10%), and deletions (10%) on the genome ℬ .

Computational efficiency was evaluated in terms of runtime and memory usage. LastZ recorded the longest runtime, taking 84.5 CPU minutes to finish, followed by wfmash (19.3 minutes) and NUCmer (2.4 minutes), while both FastGA and minimap2 completed within one minute. LastZ recorded the least peak memory usage (0.72 GB), followed by NUCmer (0.99 GB), FastGA (1.32 GB), minimap2 (1.65 GB) and wfmash (3.41 GB).

The number of alignments reported by FastGA, minimap2, LastZ, wfmash and NUCmer were 2,749, 2,724, 4,339, 1,141, and 1,974, respectively, covering 5.70 Mb, 5.36 Mb, 7.56 Mb, 15.66 Mb and 3.59 Mb sequences of the genome𝒜 . Overall, FastGA, minimap2, LastZ and NUCmer showed higher specificity than wfmash, with false aligned bases totalling 3.42 kb (0.06%), 2.53 kb (0.05%), 2.75 kb (0.04%), and 1.82 kb (0.05%), respectively, compared to wfmash’s 11.63 Mb (74.26%). Here we defined false aligned bases as alignment positions mapping outside the simulated target regions of similarity in genome 𝒜 . We also counted false positives as alignments that spanned multiple target regions or had more than 95% of their aligned bases falling outside the target region, on either genome 𝒜 or ℬ . Under this criterion, FastGA, minimap2, and NUCmer reported no false positives, LastZ reported one false positive, and wfmash reported 630 false positive alignments.

The true positive alignments were counted for every combination of region length and sequence divergence to assess the sensitivity of the genome aligners. We only considered complete alignments where their aligned bases covered at least 95% of the designated target region on both genomes. The results are depicted in Figure 7, with details showed in Supplementary Table S1. As expected, the number of target regions fully recovered by the various aligners increased with target length and decreased with sequence divergence. This trend held across all aligners except wfmash, which only produced meaningful results when the target length reached 5,000 bp. For shorter regions, most wfmash alignments were false positives spanning multiple target regions, likely due to the initial coarse-grained matching criteria used by wfmash. FastGA and minimap2 displayed very similar behaviour. Their sensitivities began to decline noticeably at sequence divergences exceeding 1%, 10%, 15%, 20%, 25%, and 30% for region lengths of 100 bp, 200 bp, 500 bp, 1000 bp, 2000 bp, and 5000 bp, respectively. FastGA was slightly less sensitive than minimap2 for smaller region lengths but more sensitive for larger regions. NUCmer had similar performance to FastGA and minimap2 at region lengths of 100 bp and 200 bp but became less sensitive than both for regions exceeding 200 bp, with the performance gap widening as region length increased. LastZ consistently demonstrated higher sensitivity than the other genome aligners. It was also the only aligner to produce reasonable alignment results at 40% sequence divergence, albeit only for regions of 2000 bp and 5000 bp.

**Figure 7:**
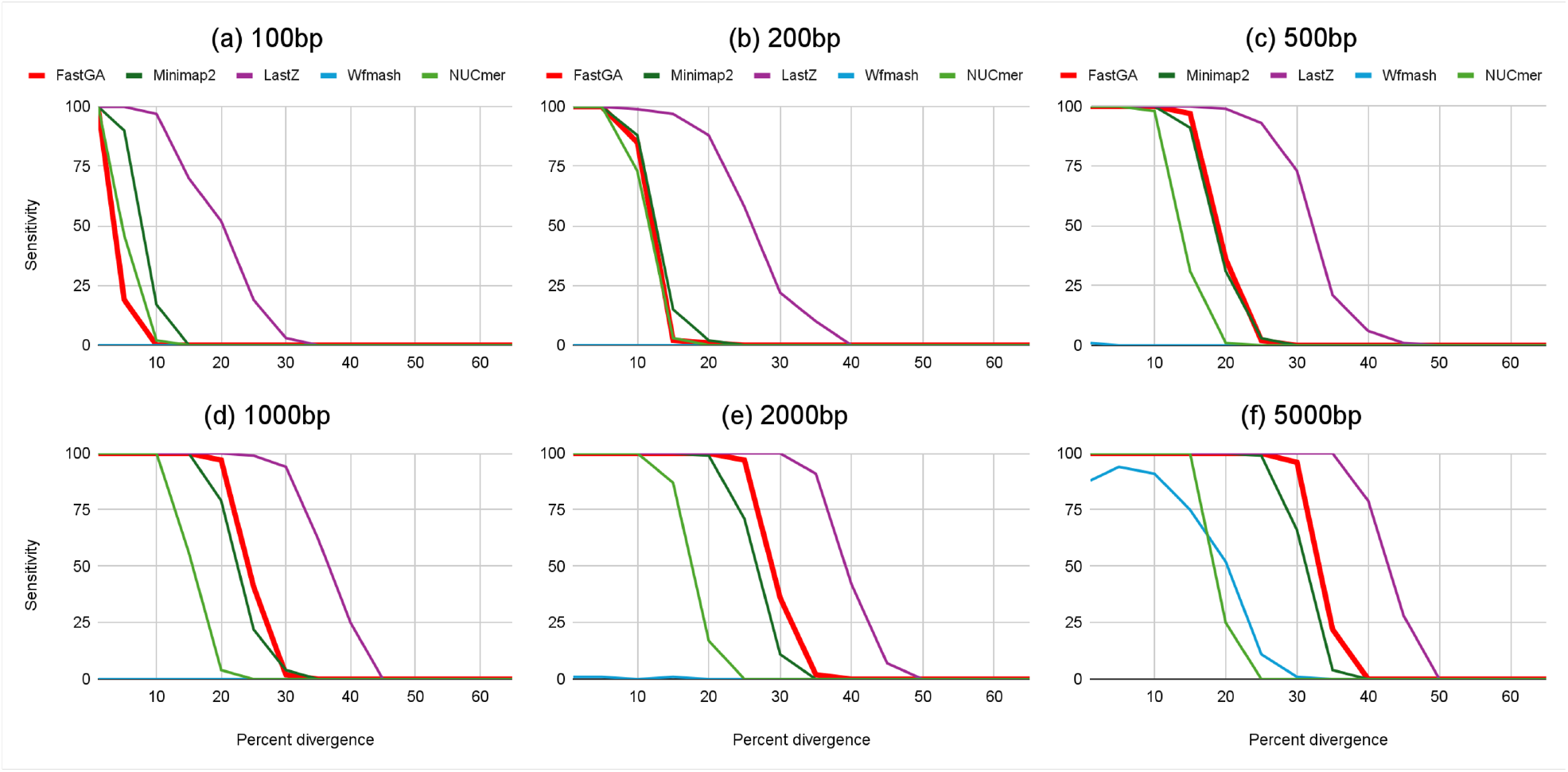
Sensitivity of genome aligners on simulated genomes. Subplots (a)-(f) correspond to target region lengths of 100 bp, 200 bp, 500 bp, 1000 bp, 2000 bp, and 5000 bp, respectively. The *x*-axis indicates the percentage of sequence divergence between the simulated genome pairs 𝒜 and ℬ , while the *y*-axis shows the number of target regions (out of 100) that were fully recovered by each genome aligner.

### 5.2 Mammalian Genomes

We selected five mammalian genomes representing a gradient of phylogenetic distances to humans and aligned them to the CHM13 “telomere-to-telomere” reference genome (3,117 Mb, GCF 009914755.1, [36]). The selected genomes, listed from closest to farthest in relation to CHM13, were: human GRCh38 (3,298 Mb, GCF 000001405.40, [37]), chimpanzee (3,178 Mb, `GCF 028858775.2,` [38]), siamang (3,263 Mb, `GCF 0288780` 55.3, [38]), pig (2,612 Mb, http://gigadb.org/dataset/102692/, [39]), and mouse (2,731 Mb, `GCA 964188535.1).`

FastGA completed genome alignments in under 71 CPU minutes for all cases, with runtime decreasing as the evolutionary distance from the human reference increased. Specifically, alignment took 70.5 minutes for human, 28.7 for chimpanzee, 26.8 for siamang, 17.4 for pig, and 16.6 for mouse. It demonstrated superior speed performance compared to all the other genome aligners, achieving speedups ranging from 29.5 × to 102.4× over minimap2, 19.1 × to 590.1 × over wfmash, 93.3 × to 843.0 × over NUCmer, and 604.8× to 2057.1× over LastZ (Figure 8a, Supplementary Table S2). FastGA also showed excellent memory efficiency, with all alignments using less than 20 GB of memory, making it feasible to run the analysis on a modern laptop. In comparison, minimap2 consumed 81 to 177 GB and wfmash consumed 84 to 105 GB of memory for these jobs. LastZ and NUCmer exhibited lower memory usages than FastGA, with a maximum consumption of only 2 GB and 5 GB, respectively (Figure 8b, Supplementary Table S2).

**Figure 8:**
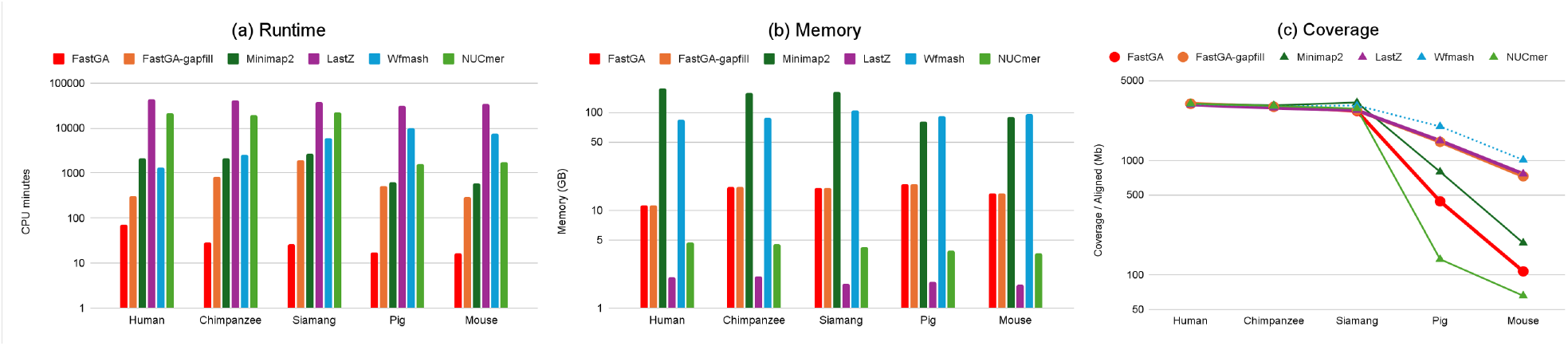
Alignment results of mammalian genomes to the CHM13 reference. Subplots (a)–(c) present the runtime, peak memory usage, and genome coverage, respectively.

The alignment results produced by different aligners vary substantially, making direct comparison of the actual alignments difficult. For instance, on the human GRCh38 genome, the number of alignments reported by FastGA, minimap2, LastZ, wfmash, and NUCmer were 579,084, 3,401, 5,628,168, 61,711 and 70,340,384, respectively. The corresponding total alignment sizes were 12,446 Mb, 3,212 Mb, 10,583 Mb, 3,124 Mb, and 94,640 Mb. We observed similar trends on the chimpanzee and siamang genomes (Supplementary Table S2). While the inherent sensitivity and specificity of genome aligners contribute to these differences, postprocessing strategies, such as alignment chaining and the handling of repetitive sequences, also play a critical role. In the GRCh38 example, minimap2 and NUCmer represent two extremes: minimap2 aggressively chains alignments and removes most repetitive matches outside primary chains, whereas NUCmer skips chaining and retains hits to repetitive regions. Despite the noticeable differences in alignment counts and total sizes, the number of GRCh38 bases covered by each aligner was very similar - 3,121 Mb, 3,134 Mb, 3,068 Mb, 3,114 Mb, and 3,135 Mb, respectively, suggesting that postprocessing more likely played the major role in the observed discrepancies. We should note that when calculating the total alignment size and the number of covered bases on the genome, we considered only the start and end positions of each alignment. As a result, gaps within alignments were included in the totals, which may affect the results - particularly for minimap2 and wfmash, which employ more aggressive chaining strategies and tend to include larger gaps.

Given the challenges of directly comparing alignment results, we used the number of genome bases covered by alignments as an indirect but meaningful metric for further assessing aligner performance (Figure 8c, Supplementary Table S2). The sensitivity of the aligners on these genomes generally aligned with what had been observed in the simulated data: FastGA and minimap2 performed similarly, both being more sensitive than NUCmer but less sensitive than LastZ. For human and chimpanzee, the genome coverages were very similar across all aligners. For siamang, FastGA, LastZ, and NUCmer produced comparable results, while minimap2 and wfmash covered 500 Mb and 300 Mb extra sequences, respectively, perhaps due to their output alignments spanning siamang-specific insertions. For the two more distantly related species, pig and mouse, differences became more pronounced, with LastZ clearly more sensitive than minimap2, FastGA and NUCmer.

Wfmash reports higher coverage than LastZ for pig and mouse, but this appear to be because of spurious false-positive matching, rather than real increased sensitivity. We can see this by considering a series of experiments in which we took a 44 Mb stretch of CHM13 chr16 (52.3-96.3 Mb) that has extended weak syntenic similarity to mouse chr8 (88.9-127.3 Mb). When directly comparing these segments, FastGA, minimap2 and NUCmer respectively mapped 3.3 Mb, 2.91 Mb and 1.1 Mb of human, while LastZ mapped 14.30 Mb and wfmash 28.4 Mb. However, when we scrambled the regions in mouse in between the FastGA matches, while LastZ dropped its coverage to only the meaningfully alignable regions with some small extensions (3.69 Mb), wfmash continued to align 22 Mb, bridging between correctly aligned regions with over 18Mb of spurious coverage. When we reverse-complemented the intervening regions LastZ recovered to 11.42 Mb, indicating that it was able to align a substantial fraction of the material in these intervals independent of consistent orientation with the more conserved flanks whereas wfmash still aligned 20.85 Mb.

In order to take advantage of the extra sensitivity of LastZ while maintaining the speed of FastGA, we explored a hybrid approach. The core idea is to use FastGA alignments as anchors and apply LastZ to fill the gaps between them. Specifically, for each pair of consistently ordered and oriented, non-overlapping FastGA alignments separated by no more than a predefined threshold (default 1 Mb), we define a bounding box using the end and start positions of the flanking alignments on the two genomes. A small overlap (default 1 kb) is allowed between the bounding box and the original alignments to facilitate seeding for LastZ. After generating these bounding boxes, we retain only the minimal ones such that no smaller ones are contained within them. LastZ is then run to align the sequence pairs corresponding to each bounding box. Finally, the alignments from FastGA and LastZ are combined to produce the final result. We present the results of this hybrid approach as ‘FastGA-gapfill’ in Figure 8 and Supplementary Table S2. As can be seen, while the sensitivity of FastGA-gapfill was comparable to LastZ, the speed was improved by a factor of 19.3 *×* to 137.5×. Although the speed gains over minimap2 and wfmash were not so strong, it remained the fastest overall, while being more sensitive than these methods.

### 5.3 Other Species

We selected twelve species sequenced by DToL ([5]) representing a wide taxonomic range, with varying genome sizes from a few hundred megabases to over 24 Gb. These include two insects from the genus *Acronicta* (*A. psi* and *A. aceris*, moths), two fishes from the genus *Thunnus* (*T. albacares* and *T. maccoyii*, tunas), two birds from the genus *Ammospiza* (*A. caudacuta* and *A. maritima*, sparrows), two reptiles from the genus *vipera* (*V. latastei* and *V. berus*, snakes), two mammals from the genus *molossus* (*M. alvarezi* and *M. nigricans*, bats), and two amphibians from the genus *Lissotriton* (*L. vulgaris* and *L. helveticus*, newts). For species from each genus, we performed two types of analyses: (1) within-species haplotype comparison, where we aligned the primary and alternative assemblies of the first species, and (2) between-species haplotype comparison, where we aligned the primary assemblies of the two different species. Further details of these genomes are provided in Supplementary Table S3.

The alignment results of FastGA are summarised in Table 1, with additional details and results from other genome aligners provided in Supplementary Table S4. FastGA completed comparisons for most of these genomes within one to two hours, using less than 12 GB memory, and consistently outperformed all other aligners in speed. For the alignment between *V. latastei* and *V. berus*, minimap2 failed under default settings and was rerun with the ‘-x asm20’ option, which is less sensitive; four LastZ jobs did not complete within the 48-hour wall-time limit, corresponding to pairwise alignments between two largest chromosomes of the genomes. FastGA was the only aligner that successfully produced alignments for the newt genomes. It took 4,611 CPU minutes (5.5 hours wall time) for the comparison between two *L. vulgaris* haplotypes, and 2,539 CPU minutes (3.2 hours wall time) for the comparison between *L. vulgaris* and *L. helveticus*, with both runs using approximately 29 GB of memory. We note that this memory requirement arose during the sorting steps to generate the genome index (GIX) files; the actual alignment process by FastGA required only around 1GB memory. Coverage fractions for all methods were comparable, except for in the comparison between the moth species, which are more distantly related, for which results were comparable to those for human to mouse.

**Table 1.** FastGA alignment results for within-species and between-species haplotype comparisons. Numbers in parentheses following the species name indicate the specific haplotypes used. The ‘CPU time’ and ‘Memory’ columns show the computational resources consumed by FastGA, while the ‘Cov A’ and ‘Cov B’ columns report the percentage of genomes A and B covered by the alignments.

| Class | Genus | Genome A | Size A (Mb) | Genome B | Size B (Mb) | CPU time (min) | Memory (GB) | Cov A (%) | Cov B (%) |
| --- | --- | --- | --- | --- | --- | --- | --- | --- | --- |
| **Insect** | *Acronicta* | *A. psi* (1) | 405 | *A. psi* (2) | 402 | 10.68 | 1.54 | 94.76 | 95.07 |
| **Fish** | *Thunnus* | *T. albacares* (1) | 792 | *T. albacares* (2) | 787 | 22.91 | 2.08 | 98.98 | 99.29 |
| **Bird** | *Ammospiza* | *A. caudacuta* (1) | 1,241 | *A. caudacuta* (2) | 1,167 | 45.40 | 2.55 | 96.44 | 99.15 |
| **Reptile** | *Vipera* | *V. latastei* (1) | 1,632 | *V. latastei* (2) | 1,549 | 153.84 | 4.03 | 93.89 | 97.96 |
| **Mammal** | *Molossus* | *M. alvarezi* (1) | 2,505 | *M. alvarezi* (2) | 2,478 | 43.51 | 10.27 | 98.84 | 99.19 |
| **Amphibian** | *Lissotriton* | *L. vulgaris* (1) | 24,226 | *L. vulgaris* (2) | 20,826 | 4,611.62 | 28.99 | 85.63 | 91.78 |
| **Insect** | *Acronicta* | *A. psi* (1) | 405 | *A. aceris* | 466 | 9.23 | 2.20 | 27.13 | 24.89 |
| **Fish** | *Thunnus* | *T. albacares* (1) | 792 | *T. maccoyii* | 782 | 21.20 | 3.87 | 98.61 | 98.84 |
| **Bird** | *Ammospiza* | *A. caudacuta* (1) | 1,241 | *A. maritima* | 1,398 | 33.15 | 3.21 | 90.40 | 89.06 |
| **Reptile** | *Vipera* | *V. latastei* (1) | 1,632 | *V. berus* | 1,695 | 70.93 | 4.23 | 91.67 | 90.53 |
| **Mammal** | *Molossus* | *M. alvarezi* (1) | 2,505 | *M. nigricans* | 2,567 | 51.24 | 11.23 | 94.77 | 93.76 |
| **Amphibian** | *Lissotriton* | *L. vulgaris* (1) | 24,226 | *L. helveticus* | 23,170 | 2,539.42 | 28.97 | 66.99 | 75.32 |

## Acknowledgements

This work was supported by the Wellcome Trust [207492 and 226458 to R.D.].

---

**Supplementary Table S1.** Alignment results of simulated genomes. The columns ‘Miss’, ‘Part’ and ‘Full’ represent the number of target regions that are missed, partially covered, or fully covered by the alignments, respectively. A target region is considered missed if it is not covered by any true positive alignments. Among the remaining targets, a region is classified as fully covered if there exists a true positive alignment covering at least 95% of its bases on both the 𝒜 and ℬ genomes; otherwise, it is considered partially covered. Rows with no hits from any aligner were omitted for simplicity.

| Length @<br>Divergence | FastGA |  |  | Minimap2 |  |  | LastZ |  |  | Wfmash |  |  | NUCmer |  |  |
| --- | --- | --- | --- | --- | --- | --- | --- | --- | --- | --- | --- | --- | --- | --- | --- |
|  | Miss | Part | Full | Miss | Part | Full | Miss | Part | Full | Miss | Part | Full | Miss | Part | Full |
| 100 @ 0.01 | 1 | 0 | 99 | 0 | 0 | 100 | 0 | 0 | 100 | 100 | 0 | 0 | 0 | 0 | 100 |
| 100 @ 0.05 | 80 | 1 | 19 | 8 | 2 | 90 | 0 | 0 | 100 | 100 | 0 | 0 | 52 | 2 | 46 |
| 100 @ 0.10 | 100 | 0 | 0 | 78 | 5 | 17 | 2 | 1 | 97 | 100 | 0 | 0 | 95 | 3 | 2 |
| 100 @ 0.15 | 100 | 0 | 0 | 100 | 0 | 0 | 22 | 8 | 70 | 100 | 0 | 0 | 100 | 0 | 0 |
| 100 @ 0.20 | 100 | 0 | 0 | 100 | 0 | 0 | 39 | 9 | 52 | 100 | 0 | 0 | 100 | 0 | 0 |
| 100 @ 0.25 | 100 | 0 | 0 | 100 | 0 | 0 | 69 | 12 | 19 | 100 | 0 | 0 | 100 | 0 | 0 |
| 100 @ 0.30 | 100 | 0 | 0 | 100 | 0 | 0 | 89 | 8 | 3 | 100 | 0 | 0 | 100 | 0 | 0 |
| 100 @ 0.35 | 100 | 0 | 0 | 100 | 0 | 0 | 98 | 2 | 0 | 100 | 0 | 0 | 100 | 0 | 0 |
| 200 @ 0.01 | 0 | 0 | 100 | 0 | 0 | 100 | 0 | 0 | 100 | 100 | 0 | 0 | 0 | 0 | 100 |
| 200 @ 0.05 | 0 | 0 | 100 | 0 | 0 | 100 | 0 | 0 | 100 | 100 | 0 | 0 | 0 | 0 | 100 |
| 200 @ 0.10 | 15 | 0 | 85 | 11 | 1 | 88 | 0 | 1 | 99 | 100 | 0 | 0 | 26 | 1 | 73 |
| 200 @ 0.15 | 98 | 0 | 2 | 83 | 2 | 15 | 0 | 3 | 97 | 100 | 0 | 0 | 97 | 0 | 3 |
| 200 @ 0.20 | 99 | 0 | 1 | 98 | 0 | 2 | 9 | 3 | 88 | 100 | 0 | 0 | 100 | 0 | 0 |
| 200 @ 0.25 | 100 | 0 | 0 | 100 | 0 | 0 | 34 | 8 | 58 | 100 | 0 | 0 | 100 | 0 | 0 |
| 200 @ 0.30 | 100 | 0 | 0 | 100 | 0 | 0 | 71 | 7 | 22 | 100 | 0 | 0 | 100 | 0 | 0 |
| 200 @ 0.35 | 100 | 0 | 0 | 100 | 0 | 0 | 87 | 3 | 10 | 100 | 0 | 0 | 100 | 0 | 0 |
| 200 @ 0.40 | 100 | 0 | 0 | 100 | 0 | 0 | 100 | 0 | 0 | 100 | 0 | 0 | 100 | 0 | 0 |
| 200 @ 0.45 | 100 | 0 | 0 | 100 | 0 | 0 | 99 | 1 | 0 | 100 | 0 | 0 | 100 | 0 | 0 |
| 500 @ 0.01 | 0 | 0 | 100 | 0 | 0 | 100 | 0 | 0 | 100 | 99 | 0 | 1 | 0 | 0 | 100 |
| 500 @ 0.05 | 0 | 0 | 100 | 0 | 0 | 100 | 0 | 0 | 100 | 100 | 0 | 0 | 0 | 0 | 100 |
| 500 @ 0.10 | 0 | 0 | 100 | 0 | 0 | 100 | 0 | 0 | 100 | 100 | 0 | 0 | 2 | 0 | 98 |
| 500 @ 0.15 | 2 | 1 | 97 | 9 | 0 | 91 | 0 | 0 | 100 | 100 | 0 | 0 | 69 | 0 | 31 |
| 500 @ 0.20 | 60 | 4 | 36 | 69 | 0 | 31 | 1 | 0 | 99 | 100 | 0 | 0 | 98 | 1 | 1 |
| 500 @ 0.25 | 97 | 1 | 2 | 96 | 1 | 3 | 7 | 0 | 93 | 100 | 0 | 0 | 100 | 0 | 0 |
| 500 @ 0.30 | 100 | 0 | 0 | 100 | 0 | 0 | 23 | 4 | 73 | 100 | 0 | 0 | 100 | 0 | 0 |
| 500 @ 0.35 | 100 | 0 | 0 | 100 | 0 | 0 | 71 | 8 | 21 | 100 | 0 | 0 | 100 | 0 | 0 |
| 500 @ 0.40 | 100 | 0 | 0 | 100 | 0 | 0 | 89 | 5 | 6 | 100 | 0 | 0 | 100 | 0 | 0 |
| 500 @ 0.45 | 100 | 0 | 0 | 100 | 0 | 0 | 97 | 2 | 1 | 100 | 0 | 0 | 100 | 0 | 0 |
| 1000 @ 0.01 | 0 | 0 | 100 | 0 | 0 | 100 | 0 | 0 | 100 | 100 | 0 | 0 | 0 | 0 | 100 |
| 1000 @ 0.05 | 0 | 0 | 100 | 0 | 0 | 100 | 0 | 0 | 100 | 100 | 0 | 0 | 0 | 0 | 100 |
| 1000 @ 0.10 | 0 | 0 | 100 | 0 | 0 | 100 | 0 | 0 | 100 | 100 | 0 | 0 | 0 | 0 | 100 |
| 1000 @ 0.15 | 0 | 0 | 100 | 0 | 0 | 100 | 0 | 0 | 100 | 100 | 0 | 0 | 43 | 1 | 56 |
| 1000 @ 0.20 | 3 | 0 | 97 | 21 | 0 | 79 | 0 | 0 | 100 | 100 | 0 | 0 | 95 | 1 | 4 |
| 1000 @ 0.25 | 59 | 0 | 41 | 78 | 0 | 22 | 1 | 0 | 99 | 100 | 0 | 0 | 99 | 1 | 0 |
| 1000 @ 0.30 | 98 | 0 | 2 | 96 | 0 | 4 | 6 | 0 | 94 | 100 | 0 | 0 | 100 | 0 | 0 |
| 1000 @ 0.35 | 100 | 0 | 0 | 100 | 0 | 0 | 35 | 3 | 62 | 100 | 0 | 0 | 100 | 0 | 0 |
| 1000 @ 0.40 | 100 | 0 | 0 | 100 | 0 | 0 | 70 | 5 | 25 | 100 | 0 | 0 | 100 | 0 | 0 |
| 1000 @ 0.45 | 100 | 0 | 0 | 100 | 0 | 0 | 96 | 4 | 0 | 100 | 0 | 0 | 100 | 0 | 0 |
| 2000 @ 0.01 | 0 | 0 | 100 | 0 | 0 | 100 | 0 | 0 | 100 | 99 | 0 | 1 | 0 | 0 | 100 |
| 2000 @ 0.05 | 0 | 0 | 100 | 0 | 0 | 100 | 0 | 0 | 100 | 99 | 0 | 1 | 0 | 0 | 100 |
| 2000 @ 0.10 | 0 | 0 | 100 | 0 | 0 | 100 | 0 | 0 | 100 | 100 | 0 | 0 | 0 | 0 | 100 |
| 2000 @ 0.15 | 0 | 0 | 100 | 0 | 0 | 100 | 0 | 0 | 100 | 99 | 0 | 1 | 13 | 0 | 87 |
| 2000 @ 0.20 | 0 | 0 | 100 | 1 | 0 | 99 | 0 | 0 | 100 | 100 | 0 | 0 | 82 | 1 | 17 |
| 2000 @ 0.25 | 3 | 0 | 97 | 29 | 0 | 71 | 0 | 0 | 100 | 100 | 0 | 0 | 98 | 2 | 0 |
| 2000 @ 0.30 | 62 | 2 | 36 | 82 | 7 | 11 | 0 | 0 | 100 | 100 | 0 | 0 | 100 | 0 | 0 |
| 2000 @ 0.35 | 98 | 0 | 2 | 97 | 3 | 0 | 9 | 0 | 91 | 100 | 0 | 0 | 100 | 0 | 0 |
| 2000 @ 0.40 | 100 | 0 | 0 | 100 | 0 | 0 | 57 | 1 | 42 | 100 | 0 | 0 | 100 | 0 | 0 |
| 2000 @ 0.45 | 100 | 0 | 0 | 100 | 0 | 0 | 91 | 2 | 7 | 100 | 0 | 0 | 100 | 0 | 0 |
| 5000 @ 0.01 | 0 | 0 | 100 | 0 | 0 | 100 | 0 | 0 | 100 | 12 | 0 | 88 | 0 | 0 | 100 |
| 5000 @ 0.05 | 0 | 0 | 100 | 0 | 0 | 100 | 0 | 0 | 100 | 6 | 0 | 94 | 0 | 0 | 100 |
| 5000 @ 0.10 | 0 | 0 | 100 | 0 | 0 | 100 | 0 | 0 | 100 | 7 | 2 | 91 | 0 | 0 | 100 |
| 5000 @ 0.15 | 0 | 0 | 100 | 0 | 0 | 100 | 0 | 0 | 100 | 17 | 8 | 75 | 0 | 0 | 100 |
| 5000 @ 0.20 | 0 | 0 | 100 | 0 | 0 | 100 | 0 | 0 | 100 | 17 | 31 | 52 | 60 | 15 | 25 |
| 5000 @ 0.25 | 0 | 0 | 100 | 1 | 0 | 99 | 0 | 0 | 100 | 45 | 44 | 11 | 97 | 3 | 0 |
| 5000 @ 0.30 | 4 | 0 | 96 | 32 | 2 | 66 | 0 | 0 | 100 | 90 | 9 | 1 | 100 | 0 | 0 |
| 5000 @ 0.35 | 74 | 4 | 22 | 87 | 9 | 4 | 0 | 0 | 100 | 99 | 1 | 0 | 100 | 0 | 0 |
| 5000 @ 0.40 | 98 | 2 | 0 | 100 | 0 | 0 | 21 | 0 | 79 | 100 | 0 | 0 | 100 | 0 | 0 |
| 5000 @ 0.45 | 100 | 0 | 0 | 100 | 0 | 0 | 70 | 2 | 28 | 100 | 0 | 0 | 100 | 0 | 0 |

**Supplementary Table S2.** Alignment results for mammalian genomes. ‘CPU time’ and ‘Memory’ measurements show the computational resources used by each aligner. ‘Alignment’ measurements present the number and total length of alignments. ‘Coverage’ measurements report the number of bases and the percentage of the genome covered by alignments.

| Measurement | Species | FastGA | FastGA-gapfill | Minimap2 | LastZ | Wfmash | NUCmer |
| --- | --- | --- | --- | --- | --- | --- | --- |
| **CPU time**<br>**(min)** | Human | 70.53 | 310.20 | 2,082.02 | 42,660.20 | 1,348.15 | 21,143.30 |
|  | Chimpanzee | 28.71 | 844.50 | 2,138.72 | 42,139.60 | 2,507.33 | 19,918.20 |
|  | Siamang | 26.82 | 1,962.50 | 2,746.12 | 37,854.90 | 6,023.16 | 22,614.40 |
|  | Pig | 17.42 | 514.90 | 625.24 | 31,485.60 | 10,280.70 | 1,624.85 |
|  | Mouse | 16.58 | 286.60 | 590.14 | 34,102.20 | 7,633.71 | 1,733.39 |
| **Memory**<br>**(GB)** | Human | 11.26 | 11.26 | 177.27 | 2.07 | 83.92 | 4.73 |
|  | Chimpanzee | 17.44 | 17.44 | 159.84 | 2.10 | 88.99 | 4.55 |
|  | Siamang | 16.94 | 16.94 | 161.97 | 1.76 | 105.26 | 4.25 |
|  | Pig | 18.64 | 18.64 | 80.70 | 1.84 | 92.82 | 3.85 |
|  | Mouse | 14.75 | 14.75 | 89.81 | 1.74 | 95.74 | 3.62 |
| **Alignment**<br>**(#)** | Human | 579,084 | 1,308,426 | 3,401 | 5,628,168 | 61,711 | 70,340,384 |
|  | Chimpanzee | 906,660 | 2,868,095 | 13,271 | 10,823,773 | 58,604 | 66,222,579 |
|  | Siamang | 1,134,864 | 6,219,092 | 49,486 | 9,720,554 | 58,753 | 77,616,820 |
|  | Pig | 301,464 | 1,341,808 | 393,480 | 1,571,272 | 99,549 | 1,539,871 |
|  | Mouse | 154,873 | 800,965 | 213,255 | 2,610,680 | 62,094 | 6,200,138 |
| **Alignment**<br>**(Mb)** | Human | 12,445.52 | 15,309.78 | 3,212.41 | 10,583.49 | 3,124.43 | 94,640.13 |
|  | Chimpanzee | 5,749.26 | 11,709.89 | 3,103.55 | 14,884.55 | 2,993.53 | 89,458.83 |
|  | Siamang | 4,325.27 | 13,520.48 | 3,363.72 | 11,433.72 | 3,036.73 | 95,675.27 |
|  | Pig | 457.69 | 2,130.13 | 805.14 | 2,190.04 | 1,998.13 | 322.46 |
|  | Mouse | 124.58 | 1,034.53 | 197.53 | 2,308.87 | 1,019.16 | 739.90 |
| **Coverage**<br>**(Mb)** | Human | 3,120.73 | 3,138.09 | 3,134.28 | 3,067.78 | 3,114.05 | 3,135.21 |
|  | Chimpanzee | 2,937.73 | 2,946.81 | 3,026.02 | 2,881.83 | 2,979.97 | 2,992.04 |
|  | Siamang | 2,712.23 | 2,747.24 | 3,210.92 | 2,743.54 | 3,017.52 | 2,840.65 |
|  | Pig | 438.70 | 1,454.24 | 795.91 | 1,491.74 | 1,988.53 | 137.00 |
|  | Mouse | 107.35 | 727.74 | 190.38 | 764.49 | 1,011.08 | 65.59 |
| **Coverage**<br>**(%)** | Human | 94.61 | 95.14 | 95.02 | 93.01 | 94.41 | 95.05 |
|  | Chimpanzee | 92.47 | 92.75 | 95.25 | 90.71 | 93.80 | 94.18 |
|  | Siamang | 83.12 | 84.19 | 98.40 | 84.08 | 92.48 | 87.06 |
|  | Pig | 16.80 | 55.68 | 30.47 | 57.11 | 76.13 | 5.24 |
|  | Mouse | 3.93 | 26.65 | 6.97 | 27.99 | 37.02 | 2.40 |

**Supplementary Table S3.** Genomes used for within-species and between-species haplotype comparisons.

| Class | Genus | Species | Haplotype | Accession NO. | Size (Mb) |
| --- | --- | --- | --- | --- | --- |
| **Insect** | *Acronicta* | *A. psi* | 1 | GCA_946251955.1 | 405 |
|  |  | *A. psi* | 2 | GCA_946251945.1 | 402 |
|  |  | *A. aceris* | - | GCA_910591435.1 | 466 |
| **Fish** | *Thunnus* | *T. albacares* | 1 | GCA_914725855.2 | 792 |
|  |  | *T. albacares* | 2 | GCA_914744365.2 | 787 |
|  |  | *T. maccoyii* | - | GCF_910596095.1 | 782 |
| **Bird** | *Ammospiza* | *A. caudacuta* | 1 | GCF_027887145.1 | 1,241 |
|  |  | *A. caudacuta* | 2 | GCA_027887135.1 | 1,167 |
|  |  | *A. maritima* | - | GCA_036010785.1 | 1,398 |
| **Reptile** | *Vipera* | *V. latastei* | 1 | GCA_024294585.1 | 1,632 |
|  |  | *V. latastei* | 2 | GCA_024294605.1 | 1,549 |
|  |  | *V. berus* | - | GCA_964194415.1 | 1,695 |
| **Mammal** | *Molossus* | *M. alvarezi* | 1 | GCA_037176705.1 | 2,505 |
|  |  | *M. alvarezi* | 2 | GCA_037157525.1 | 2,478 |
|  |  | *M. nigricans* | - | GCA_039880945.1 | 2,567 |
| **Amphibian** | *Lissotriton* | *L. vulgaris* | 1 | GCA_964263255.1 | 24,226 |
|  |  | *L. vulgaris* | 2 | GCA_964261385.1 | 20,826 |
|  |  | *L. helveticus* | - | GCA_964261635.1 | 23,170 |

**Supplementary Table S4.** Alignment results of within-species and between-species haplotype comparisons. ‘CPU time’ and ‘Memory’ measurements show the computational resources used by echo aligner. ‘Coverage’ measurements report the number of bases and the percentage of the genome covered by alignments. Missing entries marked with ‘-’ indicate that the aligner failed to produces alignments.

| Measurement | Genome A | Genome B | FastGA | Minimap2 | LastZ | Wfmash | NUCmer |
| --- | --- | --- | --- | --- | --- | --- | --- |
| CPU time<br>(min) | *A. psi* (1) | *A. psi* (2) | 10.68 | 43.86 | 1,864.66 | 129.25 | 334.19 |
|  | *T. albacares* (1) | *T. albacares* (2) | 22.91 | 52.08 | 9,676.67 | 170.29 | 517.19 |
|  | *A. caudacuta* (1) | *A. caudacuta* (2) | 45.4 | 94.4 | 17,162.20 | 193.05 | 2,591.41 |
|  | *V. latastei*(1) | *V. latastei*(2) | 153.84 | 269.06 | 28,491.20 | 606.45 | 7,090.71 |
|  | *M. alvarezi* (1) | *M. alvarezi* (2) | 43.51 | 1,371.29 | 52,755.70 | 1,518.25 | 21,515.90 |
|  | *L. vulgaris* (1) | *L. vulgaris* (2) | 4,611.62 | - | - | - | - |
|  | *A. psi* (1) | *A. aceris* | 9.23 | 58.31 | 1,853.82 | 1,646.27 | 99.56 |
|  | *T. albacares* (1) | *T. maccoyii* | 21.2 | 58.71 | 10,948.60 | 164.42 | 515.05 |
|  | *A. caudacuta* (1) | *A. maritima* | 33.15 | 436.35 | 19,780.10 | 724.7 | 4,056.05 |
|  | *V. latastei*(1) | *V. berus* | 70.93 | 1,790.07<sup>†</sup> | 87,638.40<sup>‡</sup> | 1,140.08 | 11,504.40 |
|  | *M. alvarezi* (1) | *M. nigricans* | 51.24 | 1,644.61 | 52,668.70 | 3,460.93 | 22,723.50 |
|  | *L. vulgaris* (1) | *L. helveticus* | 2,539.42 | - | - | - | - |
| Memory<br>(GB) | *A. psi* (1) | *A. psi* (2) | 1.54 | 17.89 | 0.79 | 11.76 | 1.45 |
|  | *T. albacares* (1) | *T. albacares* (2) | 2.08 | 14.86 | 0.94 | 23.32 | 1.49 |
|  | *A. caudacuta* (1) | *A. caudacuta* (2) | 2.55 | 15.72 | 1.23 | 39.63 | 2.18 |
|  | *V. latastei*(1) | *V. latastei*(2) | 4.03 | 25.9 | 2.16 | 46.97 | 3.83 |
|  | *M. alvarezi* (1) | *M. alvarezi* (2) | 10.27 | 115.46 | 3.63 | 72.31 | 4.01 |
|  | *L. vulgaris* (1) | *L. vulgaris* (2) | 28.99 | - | - | - | - |
|  | *A. psi* (1) | *A. aceris* | 2.2 | 17.19 | 0.93 | 13.85 | 1.31 |
|  | *T. albacares* (1) | *T. maccoyii* | 3.87 | 36.97 | 1.12 | 23.1 | 1.68 |
|  | *A. caudacuta* (1) | *A. maritima* | 3.21 | 47.66 | 1.86 | 39.63 | 2.82 |
|  | *V. latastei*(1) | *V. berus* | 4.23 | 194.60<sup>†</sup> | 3.09<sup>‡</sup> | 47.82 | 5.83 |
|  | *M. alvarezi* (1) | *M. nigricans* | 11.23 | 120.1 | 3.92 | 110.77 | 4.54 |
|  | *L. vulgaris* (1) | *L. helveticus* | 28.97 | - | - | - | - |
| Coverage A<br>(Mb) | *A. psi* (1) | *A. psi* (2) | 383.49 | 393.32 | 387.29 | 400.41 | 387.49 |
|  | *T. albacares* (1) | *T. albacares* (2) | 784.04 | 777.26 | 783.86 | 770.27 | 783.33 |
|  | *A. caudacuta* (1) | *A. caudacuta* (2) | 1,197.03 | 1,140.94 | 1,122.70 | 1,109.75 | 1,210.12 |
|  | *V. latastei*(1) | *V. latastei*(2) | 1,531.80 | 1,522.61 | 1,514.75 | 1,521.37 | 1,568.13 |
|  | *M. alvarezi* (1) | *M. alvarezi* (2) | 2,475.79 | 2,434.83 | 2,408.04 | 2,401.30 | 2,490.25 |
|  | *L. vulgaris* (1) | *L. vulgaris* (2) | 20,745.49 | - | - | - | - |
|  | *A. psi* (1) | *A. aceris* | 109.81 | 132.05 | 239.21 | 379.79 | 77.66 |
|  | *T. albacares* (1) | *T. maccoyii* | 781.16 | 774.32 | 782.8 | 766.29 | 778.04 |
|  | *A. caudacuta* (1) | *A. maritima* | 1,122.03 | 1,110.88 | 1,112.80 | 1,100.37 | 1,171.61 |
|  | *V. latastei*(1) | *V. berus* | 1,495.58 | 1,458.62<sup>†</sup> | 979.29<sup>‡</sup> | 1,588.62 | 1,580.56 |
|  | *M. alvarezi* (1) | *M. nigricans* | 2,373.91 | 2,356.66 | 2,369.16 | 2,355.51 | 2,424.36 |
|  | *L. vulgaris* (1) | *L. helveticus* | 16,228.07 | - | - | - | - |
| Coverage B<br>(Mb) | *A. psi* (1) | *A. psi* (2) | 382.22 | 393.15 | 384.97 | 400.62 | 387.76 |
|  | *T. albacares* (1) | *T. albacares* (2) | 781.09 | 781.14 | 781.66 | 776.16 | 782.63 |
|  | *A. caudacuta* (1) | *A. caudacuta* (2) | 1,156.86 | 1,163.65 | 1,103.34 | 1,139.26 | 1,165.17 |
|  | *V. latastei*(1) | *V. latastei*(2) | 1,518.27 | 1,542.40 | 1,497.41 | 1,540.10 | 1,544.96 |
|  | *M. alvarezi* (1) | *M. alvarezi* (2) | 2,458.68 | 2,471.95 | 2,396.78 | 2,450.11 | 2,477.27 |
|  | *L. vulgaris* (1) | *L. vulgaris* (2) | 19,114.87 | - | - | - | - |
|  | *A. psi* (1) | *A. aceris* | 116.1 | 161.48 | 259.74 | 430.24 | 90.42 |
|  | *T. albacares* (1) | *T. maccoyii* | 773.32 | 776.42 | 776.14 | 770.05 | 775.05 |
|  | *A. caudacuta* (1) | *A. maritima* | 1,244.90 | 1,362.39 | 1,183.89 | 1,302.34 | 1,387.69 |
|  | *V. latastei*(1) | *V. berus* | 1,534.53 | 1,508.07<sup>†</sup> | 997.36<sup>‡</sup> | 1,634.82 | 1,670.40 |
|  | *M. alvarezi* (1) | *M. nigricans* | 2,406.90 | 2,545.08 | 2,401.62 | 2,484.92 | 2,535.05 |
|  | *L. vulgaris* (1) | *L. helveticus* | 17,452.04 | - | - | - | - |

<sup>†</sup> Minimap2 failed with the default parameter setting and was rerun with the ‘-x asm20’ parameter.
<sup>‡</sup> Four LastZ jobs, corresponding to pairwise alignments between two largest chromosomes of *V. latastei* (1) and *V. berus*, did not complete within the 48-hour wall-time limit. These jobs were each assigned 48 hours of CPU time. Coverage calculations excluded these chromosomes.

## Footnotes

1 A MacBook Pro running macOS 10.15.7 with a 16-core M4 Max chip and 64GB of memory

2 Let the affine gap cost function be *u* + *g* where *g* is the gap length, *u* ≤ 1*/n* is a very small gap initiation charge, and *n* is the length of the sequences being compared. Then the score of an optimal unit-cost alignment would be *d* + *Gu d* + *G/n < d* + 1 where *G < n* is the number of gaps in the alignment. Therefore the best alignment under this model is an optimal unit cost alignment with the smallest number of gaps. This observation was first made by Bansho Masutani.

## References

[1] Myers, G. (2014) Efficient Local Alignment Discovery amongst Noisy Long Reads. Proc. of Algorithms in Bioinformatics WABI 2014 (Wroclaw, Poland), 52-67. Also Lecture Notes in Computer Science (Springer), vol 8701.

[2] Li, H., Handsaker, B., Wysoker, A., Fennell, T., Ruan, J., Homer, N. Marth, G., Abecasis, G., Durbin, R., 1000 Genome Project Data Processing Subgroup (2009) The Sequence Alignment/Map Format and SAMtools. Bioinformatics 25 (16), 2078–2079.

[3] Myers, E. and Durbin, R. https://www.github.com/thegenemyers/ONEcode

[4] Rhie, A., McCarthy, S.A., Fedrigo, O., Damas. J., Formenti, G., Koren, S., da Silva, M.U., … (119 more co-authors), Lewin, H.A., Howe, K., Myers, E.W., Durbin, R., Phillippy, A.M. and Jarvis, E.D. (2021) Towards Complete and Error-Free Genome Assemblies of All Vertebrate Species. Nature 592, 737–746.

[5] Blaxter, M., Mieszkowska, N., De Palma, F., Holland, D., Durbin, R.D., … and Barnes, I. (2022) Sequence Locally, Think Globally: The Darwin Tree of Life Project. PNAS 119, 4.

[6] Lewin, H.A., Richards, S., Aiden, E.L., … and Zhang, G. (2022) The Earth BioGenome Project 2020: Starting the clock. PNAS 119, 4.

[7] Li, H. and Durbin, R. (2024) Genome assembly in the telomere-to-telomere era. Nat. Rev. Genet. 25, 658–670.

[8] Lawniczak, M.K.N., Durbin, R., Flicek, P., … Zhang, G., Lewin, H.A. and Richards, S. (2022) Standards Recommendations for the Earth BioGenome Project. PNAS 119, 4.

[9] Teeling, E., Vernes, S., Davalos, L.M., Ray, D.A., Gilbert, M.T.P. and Myers, E. (2017) Bat Biology, Genomes, and the Bat1K Project: To Generate Chromosome-Level Genomes for All Living Bat Species. Annual Review of Animal Biosciences doi: 10.1146/annurev-animal-022516-022811.

[10] Altschul, S.F., Gish, W., Miller, W., Myers, E.W. and Lipman, D.J. (1990) Basic local alignment search tool. Journal of Molecular Biology. J. of Molecular Biology 215, 403–410.

[11] Paten B, Earl D, Nguyen N, Diekhans M, Zerbino D, and Haussler D. (2011) Cactus: Algorithms for genome multiple sequence alignment. Genome Research 21, 9, 1512-28.

[12] Armstrong, J., Hickey, G., Diekhans, M. … Zhang, G., and Paten, B. (2020) Progressive Cactus is a multiple-genome aligner for the thousand-genome era. Nature 587, 246–251.

[13] Sierra, P. and Durbin, R. (2024) Identification of transposable element families from pangenome polymorphisms. Mob DNA 15, 13.

[14] Schwartz, S., Zheng, Z., Frazer, K.A., Smit, A., Riemer, C., Bouck, J., Gibbs, R., Hardison, R.C., and Miller, W. (2000) PipMaker - A Web Server for Aligning Two Genomic DNA Sequences. Genome Research 10, 577–586.

[15] Schwartz, S., Kent, J.W., Smit, A., Zheng, Z., Baertsch, R., Hardison, R.C., Haussler, D., and Miller, W. (2003) Human-Mouse Alignments with BLASTZ. Genome Research 13, 103–107.

[16] Harris, R.S. (2007) Improved pairwise alignment of genomic DNA. Ph.D. Thesis, The Pennsylvania State University.

[17] Ma, B., Tromp, J. and Li, M. (2002) PatternHunter: faster and more sensitive homology search Bioinformatics 18, 440–445

[18] Marcals, G., Delcher, A.L., Phillipy, A.M., Coston, R., Salzberg, S.L. and Aleksey, Z. (2018) MUMmer4: A fast and versatile genome alignment system. PLoS Computational Biology 14 (1), e1005944.

[19] Heng, L. (2018) Minimap2: pairwise alignment for nucleotide sequences. Bioinformatics 34 (18), 3094–3100.

[20] Heng, L. (2018) New strategies to improve minimap2 alignment accuracy. Bioinformatics 37 (23), 4572–4574.

[21] Roberts, M., Hayes, W., Hunt, B.R., Mount, S.M., Yorke, J.A. (2004) Reducing storage requirements for biological sequence comparison. Bioinformatics 20 (18), 3363–3369.

[22] Garrison, E. and Guarracino, A. (2022) Unbiased Pangenome Graphs. Bioinformatics 39 (1), 1–7.

[23] Jain, C., Koren S., Dilthey, A., Phillippy, A.M., and Aluru, S. (2018) A Fast Adaptive Algorithm for Computing Whole-Genome Homology Maps Bioinformatics 34 (ECCB 2018), i748–i756.

[24] Marco-Sola S., Moure, J.C., Moreto, M., and Espinosa, A. (2020) Fast gap-affine pairwise alignment using the wavefront algorithm. Bioinformatics 37 (4), 456–463.

[25] Edgar, R. (2021) Syncmers Are More Sensitive than Minimizers for Selecting Conserved k-mers in Biological Sequences. PeerJ 9, e10805, doi: 10.7717/peerj.10805.

[26] Kielbasa, S.M., Wan, R., Sato, K., Horton, P. and Frith, M.C. (2011) Adaptive Seeds Tame Genomic Sequence Comparison. Genome Research 21, 487–493.

[27] Kokot, M., Deorowicz, S., and Debudaj-Grabsyz, A. (2017) Sorting Data on Ultra-Large Scale with RADULS. Proc. of Beyond Databases, Architecture and Structures (vol. 716, Springer), 235–245.

[28] Satish, N., Kim, C., Chugani, J., Nguyen, A.D., Lee, V.W., Kim, D. and Dubey, P. (2010) Fast Sort on CPUs and GPUs: A Case for Bandwidth Oblivious SIMD Sort. Proc. of ACM SIGMOD Conf. on Management of Data, 351–362.

[29] Cho, M., Brand, D., Bordawekar, R., Finkler, U., Kulandaisamy, V. and Puri, R. (2015) Paradis: an efficient parallel algorithm for in-place radix sort. Proc. of the VLDB Endowment 8,12, 1518–1529.

[30] Abouelhoda, M.I., Kurtz, S., Ohlebusch, E. (2004) Replacing Suffix Trees with Enhanced Suffix Arrays. J. of Discrete Algorithms 2, 53–86.

[31] Myers, E.W. (1986) An O(ND) difference algorithm and its variations. Algorithmica 1, 251–266.

[32] Ukkonen, E. (1985) Algorithms for Approximate String Matching. Information and Control 64, 100–118.

[33] Wu, S., Myers, E., Manber, U., and Miller, W. (1990) An O(NP) Sequence Comparison Algorithm. Information Processing Letters 35 (6), 317–323.

[34] Hadlock, F. (1988) Minimum Detour Methods for String or Sequence Comparison. Congressus Numerantium 61, 263–274.

[35] Martiniano, R., De Sanctis, Hallast, P. and Durbin, R. (2022) Placing Ancient DNA Sequences into Reference Phylogenies. Mol Biol Evol 39:msac017.

[36] Nurk, S., Koren, S., Rhie, A., Rautiainen, M., Bzikadze, A.V., Mikheenko, A., Vollger, M.R., Altemose, N., Uralsky, L., Gershman, A. and Aganezov, S. (2022) The complete sequence of a human genome. Science 376(6588), 44–53.

[37] Schneider, V.A., Graves-Lindsay, T., Howe, K., Bouk, N., Chen, H.C., Kitts, P.A., Murphy, T.D., Pruitt, K.D., Thibaud-Nissen, F., Albracht, D. and Fulton, R.S. (2017) Evaluation of GRCh38 and de novo haploid genome assemblies demonstrates the enduring quality of the reference assembly. Genome Research 27(5), 849–864.

[38] Yoo, D., Rhie, A., Hebbar, P., Antonacci, F., Logsdon, G.A., Solar, S.J., Antipov, D., Pickett, B.D., Safonova, Y., Montinaro, F. and Luo, Y. (2025) Complete sequencing of ape genomes. Nature 1–18.

[39] Cao, C., Miao, J., Xie, Q., Sun, J., Cheng, H., Zhang, Z., Wu, F., Liu, S., Ye, X., Gong, H. and Zhang, Z. (2025) A near telomere-to-telomere genome assembly of the Jinhua pig: enabling more accurate genetic research. GigaScience 14, p.giaf048.

